# Stretch versus shortening contractions subsequently decrease versus increase neural drive to the human tibialis anterior

**DOI:** 10.64898/2026.03.13.710530

**Authors:** Brent James Raiteri, Karla Friederike Bosse, Marta Boccardo, Alain Charles Vandal, Daniel Hahn

## Abstract

EMG-based muscle force predictions are often inaccurate following active muscle stretch or shortening because of residual force enhancement (rFE) or depression (rFD), respectively, which can alter the neural drive to a muscle. However, the extent of neural drive modulation due to rFE or rFD remains unknown, making it difficult to correct EMG-based force predictions. Therefore, seventeen participants performed dorsiflexion contractions at 20 and 40% of maximum voluntary torque (MVT) in three conditions: stretch-hold, shortening-hold, and fixed-end reference (REF) conditions. The ankle dorsiflexion torques and angles were matched using dynamometry to the REF condition over a 10-s steady state following a 1-s 25° stretch or shortening, during which we recorded and decomposed tibialis anterior individual motor unit action potentials from high-density surface EMG recordings to gain insights into neural drive. Normalized EMG amplitudes were 2% lower following stretch and 1 or 3% higher following shortening relative to REF at 20 versus 40% MVT (*p*≤.008), respectively. Discharge rates (DRs) from 19 matched motor units per person on average obtained via DEMUSE and MUedit were similar (*p*=.871). Following stretch and shortening, DRs were ∼1 Hz lower (*p*≤.004) and 0 (*p*=.966) to 1 Hz higher relative to REF (*p*=.003), respectively. More unique motor units were also detected following shortening versus REF and in REF versus following stretch. These findings indicate that to account for rFE or rFD, neural drive is respectively decreased or increased via reduced or additional motor unit recruitment and DR modulation, with a contraction-intensity specific discharge rate modulation following active shortening.

## INTRODUCTION

One of the holy grails of biomechanics is to accurately predict muscle force, as direct measurements of muscle force are currently impossible. Accurate muscle force predictions are crucial for estimating tissue and joint loading (Navacchia *et al*., 2016), as well as for informing injury prevention and rehabilitation strategies. However, accurate predictions are challenging as information is required on joint kinematics, activation dynamics and muscle lengths and their rates of change (Zajac, 1989; Pandy *et al*., 1990). Even for a relatively simple scenario where activation is fixed, steady-state muscle force predictions are often inaccurate, with force being higher than predicted following active muscle stretch or lower than predicted following active muscle shortening (Edman *et al*., 1978; Granzier & Pollack, 1989; Leonard *et al*., 2010; Trecarten *et al*., 2015). These prediction inaccuracies arise because of two phenomena known as residual force enhancement (rFE) and residual force depression (rFD) (Abbott & Aubert, 1952). rFE and rFD respectively describe the enhanced or depressed steady-state force following active muscle stretch or shortening relative to the force attained during fixed-end reference contractions at a matched final muscle length and activation level.

rFE and rFD are typically observed in activation-matched (*ex vivo*) or EMG-amplitude-matched (*in vivo*) experiments (Abbott & Aubert, 1952; Herzog *et al*., 1998; Pinniger & Cresswell, 2007), with reported amplitudes from 7-27% and 6-24%, respectively. If force or torque is matched rather than activation/activity level, there is thus the potential for either rFE or rFD to alter the underlying neural drive for a given muscle force, which has implications for accurately predicting muscle force based on EMG measurements (Lloyd & Besier, 2003). Previous *in vivo* results from force/torque-matched experiments indicate lower (5-44%) or higher (9-48%) EMG amplitudes in the steady state following active muscle-tendon unit stretch or shortening, respectively (Oskouei & Herzog, 2005; Rousanoglou *et al*., 2007; Altenburg *et al*., 2008; Paquin & Power, 2018; Dalton *et al*., 2018; Jakobi *et al*., 2020). Consequently, neural drive might similarly change following active stretch or shortening. However, neural drive changes cannot be inferred from EMG amplitude changes derived from conventional bipolar surface EMG because of the effects of peripheral muscle properties on EMG amplitude estimations (Dimitrov *et al*., 2008; Martinez-Valdes *et al*., 2018; Dideriksen & Farina, 2019). Rather, to accurately assess neural drive, the individual discharge patterns of the active alpha motoneurons need to be assessed (Caillet *et al*., 2024).

Only a couple of studies have used intramuscular EMG to assess neural drive changes following active muscle-tendon unit stretch or shortening. Altenburg and colleagues (2008) found that vastus lateralis motor unit discharge rates were respectively similar (mean (SD): -0.1 (0.1) Hz; -1 (8%)) or higher (0.7 (0.4) Hz; 11 (9)%) following stretch or shortening, whereas EMG amplitudes were lower (-16 (12)%) or higher (9 (6)% to 19 (14)%) following respective stretch or shortening relative to torque-matched (4-50% of maximum) fixed-end reference contractions. More recently, Jakobi and colleagues (2020) found that tibialis anterior (TA) motor unit discharge rates were lower (mean range: -2.1 to -3.6 Hz; - 19 to -27%) following stretch (shortening was not studied) relative to torque-matched (10 and 20% of maximum) fixed-end reference contractions. Additionally, these authors reported lower TA EMG amplitudes and reduced torque steadiness following active muscle-tendon unit stretch versus fixed-end reference contractions, which was attributed to a net decline in the overall activity of the motoneuron pool (-24 (16) to -44 (24)%) and not to increased discharge rate variability (Jakobi *et al*., 2020). However, only a relatively limited number of motor units were studied per person (≤7), which is a general limitation of intramuscular EMG due its high spatial selectivity (McGill & Dorfman, 1985). Further studies are thus needed to assess neural drive changes based on a relatively larger number of active motor units to better understand the behaviour of the active motoneuron pool following both stretch and shortening.

One way to non-invasively identify discharge patterns from a larger number of individual motor units is through the use of decomposition algorithms alongside high-density surface EMG (HDsEMG) measurements (Holobar & Zazula, 2004, 2007*a*). Non-invasive HDsEMG decomposition has comparable accuracy to intramuscular EMG decomposition for identifying individual motor unit discharge times (Holobar *et al*., 2009, 2010; Marateb *et al*., 2011; Negro *et al*., 2016) and could provide new insight into neural drive changes following contraction-history-dependent changes to a muscle’s force capacity. Therefore, with HDsEMG decomposition, we aimed to determine the extent of neural drive modulation from a larger population of active motor units than studied previously following muscle-tendon unit stretch or shortening contractions compared with torque- and joint-angle-matched fixed-end reference contractions. To increase the likelihood of inducing rFE or rFD, we used an active preload phase prior to muscle-tendon unit stretch or shortening to ensure active muscle stretch or shortening during muscle-tendon unit displacement (Raiteri *et al*., 2024). We chose to study the human tibialis anterior (TA) muscle because accurate single motor unit discharges can be decomposed up to 70% of maximal voluntary contraction for this muscle (Holobar *et al*., 2014), and because the TA contributes to roughly half of the voluntary dorsiflexion torque based on its physiological cross-sectional area (Brand *et al*., 1986). Additionally, the TA undergoes both active stretch and shortening contractions during walking (Maharaj *et al*., 2019), so understanding how active length changes affect TA’s neural drive in a carefully-controlled experiment could inform us about potential neural drive modulations during everyday locomotion.

We hypothesized that i) normalized EMG amplitudes and discharge rates would be respectively lower or higher during the steady state following active muscle-tendon unit stretch or shortening compared with fixed-end reference contractions. We also expected that ii) the decrease in neural drive following stretch would be similar among contraction intensities, whereas the increase in neural drive following shortening would be larger at 40 versus 20% of maximal voluntary torque. This is because rFE, in relative terms, appears to be independent of contraction intensity (Oskouei & Herzog, 2006; Seiberl *et al*., 2012), whereas relative rFD appears to increase with increasing contraction intensity (De Ruiter *et al*., 1998; Raiteri *et al*., 2024). In addition, we expected that iii) changes in EMG amplitudes and motor unit discharge rates would be strongly (but not perfectly) and positively correlated. Lastly, based on previous findings (Jakobi *et al*., 2020), we hypothesized that iv) torque variability but not motor unit discharge rate variability would be higher following stretch compared with following shortening and during fixed-end reference contractions.

## METHODS

### Ethical approval

All experimental procedures were approved by the Faculty of Sport Science’s ethics committee at Ruhr University Bochum (EKS V 09/2022) and were conducted in accordance with the Declaration of Helsinki, aside from database registration. Free written informed consent was obtained prior to the first testing session.

### Sample size calculation

A minimum sample size of sixteen was calculated with G*Power (v3.9.1.7, RRID:SCR_01372637; Faul *et al*., 2009) to achieve at least 95% power to detect a minimum effect size of interest (*d*_z_) of 1 between two conditions of interest with a two-tailed alpha level of 5%. Consequently, eighteen healthy and recreationally active participants (seven women) were tested to allow for two dropouts.

### Participants

Participants were free of pain, injury, and fatigue within their lower limbs at the time of testing. Each participant completed one familiarisation and one testing session, which were separated by two to seven days (4.5 (1.8) days). Only one female participant was considered a dropout because they were unable to accurately match the torque traces in the three experimental conditions described below, which resulted in seventeen usable datasets (n=17, age: 25 (3) yr [min-max: 22-31 yr], height: 1.77 (0.06) m [min-max: 1.65-1.85 m]; mass: 74 (9) kg [min-max: 58-90 kg], six women).

### Experimental setup

The experimental setup was almost identical to that already described and shown in Figure 1 of Raiteri et al. (2023) and only a brief description will be given here. Participants sat in a reclined posture with their right hip and knee at ∼110° and ∼90° flexion, respectively, and the sole of their right foot secured to the dorsi/plantar flexion attachment of a motorized dynamometer (IsoMed2000, D&R Ferstl GmbH, Hemau, Germany). A dorsiflexion attachment over the metatarsals ensured that the participant’s forefoot maintained constant contact with the dynamometer footplate during voluntary contractions of their right dorsiflexors. This dorsiflexion attachment also limited ankle joint rotation and force contributions from the toe extensors to the measured net ankle dorsiflexion torque due to its rigidity and location over the foot. Net ankle joint torque and crank arm angle were sampled at 2 kHz by the dynamometer and digitally stored via a Spike2 data collection system (16-bit Power1401-3 and 64-bit 8.23 Spike2 version; Cambridge Electronic Design Ltd., Cambridge, United Kingdom).

**Figure 1.**
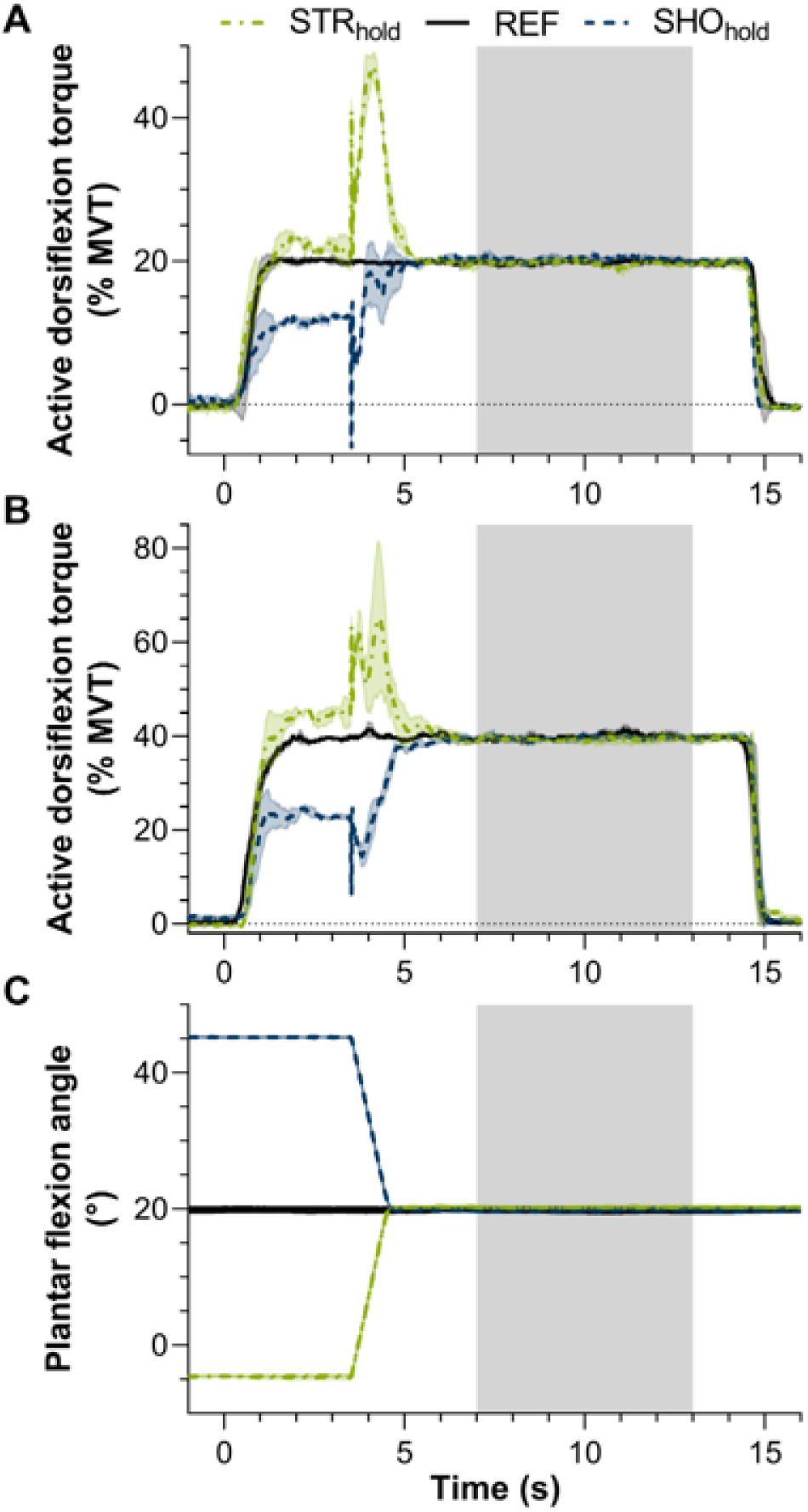
Overview of the tested contraction intensities (A: 20% of maximal voluntary torque, MVT; B: 40% MVT), and (C) experimental conditions; the mean trace of multiple trials per condition from one participant are shown, with shaded areas around the mean indicating the standard deviation and the grey shaded area indicating the 6-s steady-state phase. Participants were required to match contraction intensities at (A) 20 and (B) 40% MVT (normalized to 20° plantar flexion) in (C) three conditions: stretch-hold (STR_hold_) contractions, fixed-end reference (REF) contractions, and shortening-hold contractions (SHO_hold_). For STR_hold_ and SHO_hold_, the torque in A and B prior to the rotation in C differs from 20 and 40% MVT as the joint-angle-specific normalized torque was matched, whereas here the torque traces have been normalized to the MVT produced at 20° plantar flexion; after the rotation, the joint-angle-specific normalized torque at 20° plantar flexion was then matched.

Simultaneously, 64 HDsEMG channels were sampled in monopolar mode from one two-dimensional grid (13 rows × 5 columns of electrodes with one missing electrode in the top corner, 3 mm recording diameter, 8 mm interelectrode distance; GR08MM1305, OT Bioelettronica, Italy) over the mid-belly of the TA muscle at 2 kHz by Sessantaquattro hardware (16-bit resolution with a 10.5 Hz high-pass first-order recursive exponentially-weighted moving average firmware filter and a 500 Hz low-pass third-order digital sinc filter also with frequency notches at 2000 Hz and its harmonics, 1 V/V gain) and stored via OT BioLab+ software (OT Bioelettronica, Italy). Antagonistic muscle activity from the soleus muscle was not assessed based on previous findings of similar EMG amplitudes following active muscle stretch or shortening relative to fixed-end reference contractions at 20, 40, 60 and 80% of maximum (Paquin & Power, 2018). The reference electrode was located above the right fibular head (8 mm recording diameter, Ag/AgCl, Kendall, Mansfield, Massachusetts, United States). Electrode and grid placement occurred after the skin was shaved (Teqler, Wecker, Luxembourg), exfoliated (Nuprep, Weaver and Company, Aurora, United States) and cleaned (Sterillium, HARTMANN, Heidenheim, Germany), and after the holes of a disposable bioadhesive foam layer, which was positioned between the grid and skin, were filled with conductive cream to enable skin-electrode contact (AC Cream, Spes Medica, Genoa, Italy). EMG data was later synchronized with torque and angle data via a 5V TTL signal with rising and falling phases at the start and end of the recording, respectively, that was output by the Spike2 data collection system and recorded at 2 kHz by both systems.

### Experimental protocol

Participants were familiarized with the different experimental conditions during a practice session to improve torque matching during the actual testing session. In both sessions, participants received real-time visual feedback of their net ankle joint torque (i.e. 0.01-s moving average) via a screen positioned in front of them. The warm-up consisted of five fixed-end dorsiflexion contractions (1-s hold, 1-s rest) at ∼80% of the participant’s perceived maximum at 20° plantar flexion (note that 0° plantar flexion represents a 90° angle between the tibia and sole of the foot). Participants then performed maximal voluntary contractions (MVCs, 3-s to 5-s hold) with their right dorsiflexors at -5°, 20° and 45° plantar flexion in a randomized order under standardized verbal encouragement from the investigator. If the difference in peak-to-peak torque between two maximal-effort trials at the same joint angle was >5%, additional maximal-effort trials were performed until this difference was ≤5%, and a three-minute break was provided after each maximal-effort trial. The trial with the highest peak-to-peak torque (maximal voluntary torque, MVT) that was within 5% of another trial at each joint angle was then taken so that participants could match joint-angle specific contraction intensities of 20 and 40% MVT at -5°, 20° and 45° plantar flexion.

Participants subsequently performed three conditions at 20 and 40% MVT. The order of the conditions and the order of the contraction intensities was block randomized, such that all conditions at one contraction intensity were performed before the other contraction intensity, and trials from one condition were repeated before trials of another condition started. The three conditions included two dynamic conditions and one fixed-end reference (REF) condition, and all conditions had a common final ankle joint angle of 20° plantar flexion. The REF condition involved a 1-s ramp up to 20 or 40% MVT and a 13-s hold phase (Fig. 1). The stretch-hold (STR_hold_) and shortening-hold (SHO_hold_) conditions involved a 1-s ramp up to 20 or 40% MVT and 2-s of pre-activation before the ankle dorsiflexor muscle-tendon units were stretched or shortened. The stretch or shortening subsequently occurred over 1-s (i.e. 25°·s^-1^) from an ankle joint angle of -5° or 45° plantar flexion to 20° plantar flexion. Following the dynamic phase, a 10-s hold phase at 20 or 40% MVT was maintained at 20° plantar flexion.

In each condition, participants attempted to keep their net ankle joint torque between two traces that were located 5% above and below the desired contraction intensity. However, participants were instructed to maintain the same perceived effort during the 1-s stretch or shortening phase due to the difficulty of matching torque during the dynamic phase (Fig. 1). Participants practised each condition once at 20 or 40% MVT before repeating four trials of each condition at the same contraction intensity, resulting in 24 test trials in total. Each trial was followed by a one-minute break.

### Data processing and analysis

#### Synchronization

The first data processing step involved using a custom-written script to export individual trials within each Spike2 recording as separate files with a .mat extension. The recorded digital signals from each trial were then cropped in MATLAB (64-bit R2022a version, MathWorks, Natick, Massachusetts, United States) between a common start and end time using a voltage-crossing threshold of 95% of the maximum 5-V TTL signal recorded. The cropped torque and angle signals of each trial were then combined with the corresponding HDsEMG signals, which were imported into MATLAB using [OpenOTBfilesConvFact.m](https://github.com/OTBioelettronica/OTB-Matlab/tree/main/MATLAB%20Open%20and%20Processing%20OTBFiles/OpenOTBFiles) and cropped between a common start and end time using a voltage-crossing threshold of 10 times the minimum TTL signal recorded. Different voltage-crossing thresholds were used to synchronize the data because of different temporal lags (0.5 ms for Spike2 and 33.2 (0.1) ms for OTB), voltage input ranges (10 V for Spike2 and 6.6 V for OTB) and amplitude inversions between systems (positive for Spike2 and negative for OTB).

#### Filtering and normalization

As in previous work (Raiteri *et al*., 2024), recorded torque (20 Hz) and crank arm angle (6 Hz) data were low-pass filtered using dual-pass second-order Butterworth filters that were corrected for two passes (Winter, 2009). A steady-state passive net ankle joint torque-angle fit was then constructed to estimate active torque, which involved fitting a second-order polynomial to the mean filtered ankle joint angle and net ankle joint torque data calculated over 1 s at the start of each trial and over 0.25- s at the end of each trial when participants were instructed to relax. Active torques during each trial were subsequently estimated by evaluating the polynomial fit at the crank arm angles recorded during the trial and subtracting these values from the recorded net ankle joint torques. TA’s EMG amplitudes during each trial were calculated by bandpass filtering (20 to 400 Hz) single-differential signals with a fourth-order Butterworth filter and then smoothing these filtered and DC-removed signals with a centred moving (250-ms or 99.8% overlap) 250.5-ms root mean square (RMS) amplitude calculation. The single-differential signals were calculated from consecutive electrode pairs in the grid’s longitudinal direction, leading to 59 signals for TA EMG amplitude (four of the five columns with 12 bipolar electrode pairs and one column with 11 pairs). Each single-differential signal was later normalized by its respective value from the MVC with the highest active torque at 20° plantar flexion to determine normalized TA EMG amplitudes. To calculate the 59 normalization values, the time instant of the maximal RMS amplitude of each signal was first calculated, and then the centred mean RMS amplitude over a 500-ms window from this index was determined.

#### Time averaging

Torque steadiness was calculated as the coefficient of variation (CV) in active torque during the steady state. The steady state was 6 s in duration and defined from 7 s after contraction onset to 1 s before the end of the hold phase (Fig. 1). The means for the outcome variables of interest (i.e. active torque, torque steadiness, normalized TA EMG amplitude, TA discharge rate variability; except for TA discharge rate) were calculated for each valid trial during this steady state. Individual motor unit discharge rates were instead calculated as the median discharge rate during the steady state of each valid trial to avoid outliers from biasing the central tendency. A trial was deemed valid if there was less than a 1.0 Nm active torque difference between the first second and last second of the steady-state period.

#### Motor unit decomposition

Only the steady-state phase (grey area in Fig. 1) of the valid trials was decomposed to improve accuracy by minimising temporal and spatial changes to the waveforms of motor unit actional potentials. Trials were excluded from analysis if their mean normalized EMG amplitude fell outside one median absolute deviation of the median from valid trials in that condition to avoid outliers from biasing the results, which resulted in 2 to 5 valid trials per condition. The valid trials from each condition were then concatenated together at each contraction intensity to allow the same motor units to be tracked among conditions. Subsequently, the concatenated conditions for decomposition were reformatted (.mat) to be read by the decomposition software of interest and included 6 to 11 valid trials of 36 to 66 s in duration. Notably, the concatenated trials for decomposition were 1 s longer (i.e. 37 to 67 s) in duration because the respective start and end of the first and last concatenated trials needed to be padded with 0.5 s of real data to avoid omission during EMG preprocessing by the decomposition software. These files were then imported into the decomposition software and the raw HDsEMG signals were visually inspected to remove channels with motion artifacts or bad signal-to-noise ratios (1 channel was removed from two participants at 20% MVT, and 1 channel was removed from 1 participant at 40% MVT). The remaining channels were then pre-processed according to the defaults of DEMUSE (Holobar & Zazula, 2007*b*, 2007*a*) and MUedit (Avrillon *et al*., 2024) (see supporting information for details) and decomposed into individual motor unit action potentials using blind source separation (note that the source code of MUedit was edited to remove manual cropping and ensure automatic selection of the full file duration). Contraction intensities were decomposed separately to avoid biasing the decomposition results to higher-threshold motor units active at 40% but not 20% MVT.

DEMUSE (Holobar & Zazula, 2007*b*, 2007*a*) and MUedit (Avrillon *et al*., 2024) were both used to enhance the robustness of the decomposition results. Their approaches are similar but not identical, with DEMUSE (v5.01; University of Maribor, Slovenia) using a gradient convolution kernel compensation (gCKC) algorithm (Holobar & Zazula, 2007*b*) and MUedit using the method developed by Negro et al. (2016) that incorporates fast independent component analysis (Chen & Zhou, 2016) and gCKC-based source refinement. Nevertheless, both approaches aim to identify a set of filters that distinguish between surface-recorded motor unit action potentials based on their shape and location to isolate motor unit discharge patterns. Each approach has been validated previously up to 70 (Holobar *et al*., 2014) or 90% MVT (Negro *et al*., 2016) by identifying the same motor unit from intramuscular EMG signals and blind source separation of the HDsEMG signals recorded from TA. We opted for an extension factor of ∼16 (i.e. 1000 signals) for all decompositions based on the recommendation to reach 1000 signals to maximise the number of identified motor units (Negro *et al*., 2016). Other decomposition parameters for DEMUSE and MUedit can be found in the supporting information.

#### Decomposition editing

The decomposition outputs from DEMUSE were manually edited by one expert (MB) and the motor unit discharge patterns recalculated based on previous guidelines (Del Vecchio *et al*., 2020; Hug *et al*., 2021). In line with these guidelines, there was no subjective selection of motor unit discharges in the final decomposition output. Additionally, duplicate motor units in the final output that shared more than 30% of their identified discharge times (within a time interval of 0.5 ms) were removed with the lower PNR motor unit being excluded (Holobar *et al*., 2010).

The decomposition outputs from MUedit were also initially manually edited by one operator (BJR) and recalculated based on the same guidelines. However, we found that an automatic approach with a median analysis of motor unit discharge rates resulted in negligible differences in discharge rate compared with the mean motor unit discharge rate determined following manual editing. Consequently, we employed this automatic editing approach for MUedit decompositions due to its objectivity and time efficiency. However, this automatic editing approach did not work for DEMUSE decompositions because the CV-based exclusion led to the removal of most motor units (i.e. 0 to 4 remaining, see below). The automatic editing approach first involved removing duplicate motor units based on the same criterion (i.e. 30% of shared timings) as above (Holobar *et al*., 2010). Then, we removed the automatically-detected-and-edited motor units from MUedit with a: 1) pulse-to-noise ratio (PNR) of less than 28 dB, which corresponds to an accuracy of less than 85% (Holobar *et al*., 2014), and; 2) CV in discharge rate of 40% or above (Kalc *et al*., 2023). This CV-based exclusion worked for MUedit decompositions because unlike DEMUSE, MUedit refines separation vectors until the CV in inter-spike intervals is minimised or below 30%. As we opted for a PNR metric that does not rely on the regularity of a motor unit discharge pattern to estimate the accuracy of discharge identification, we also opted to use a CV threshold to remove mixed clustered discharge patterns (Negro *et al*., 2016). The motor unit discharge patterns were then visually inspected, but manual editing (i.e. adding missing and removing falsely identified discharge times) was deemed unnecessary because the median discharge rate of each motor unit per condition was minimally affected by this process. Interested readers can find the agreement between the manually edited DEMUSE decomposition results and the automatically edited MUedit decomposition results in the supporting information.

### Statistics

A two-tailed alpha level was set at 5% and statistical analysis was performed using R (v4.2.2; R Core Team, 2022). We fitted linear mixed-effects models (estimated using maximum likelihood and the nloptwrap optimizer from the lme4 package (Bates *et al*., 2015)) to investigate the fixed (i.e. fixed slopes, but random intercepts) and random (i.e. random slopes and intercepts) contraction intensity and condition effects on the outcome variables of interest (i.e. active torque, torque steadiness, normalized EMG amplitude, discharge rate and discharge rate variability). Mixed-effects models were primarily used to appropriately handle the varying number of motor units per person. To determine which model better predicted each outcome variable of interest, we systematically compared models with a contraction intensity × condition interaction using the flexplot package (Fife, 2022). For each outcome variable, the random-effects model without random trial effects nested under random participant effects resulted in a better fit based on lower values for the Akaike Information Criterion and Bayesian Information Criterion and a higher Bayes factor (see supporting information). No interactions were included to fit active torque, torque steadiness or discharge rate variability, whereas a random contraction intensity × condition interaction effect was included to fit normalized EMG amplitude and discharge rate. For normalized EMG amplitude, we used a fixed contraction intensity × condition interaction effect with random channel effects nested under random participant effects. For both motor unit discharge rate and discharge rate variability, we also investigated the random effect of decomposition algorithm, with random motor unit effects nested under random participant effects. Sandwich variance estimation-based standard errors (type “CR2” due to the small number of clusters) and *p*-values were calculated using the clubSandwich package (Pustejovsky *et al*., 2025) and reported below as these do not assume normality of the residuals or homoscedasticity between contraction intensities. We used the partR2 package (Stoffel *et al*., 2021) to partition the total proportion of variance explained by fixed effects (inclusive R^2^) in each model. To decompose a significant contraction intensity × condition interaction, we conducted three post-hoc pairwise comparisons at each contraction intensity using the emmeans package (Lenth *et al*., 2025) and calculated Holm-adjusted *p* values to control the family-wise error rate at 5%. Degrees of freedom were estimated using the Satterthwaite approximation with the lmerTest package (Kuznetsova *et al*., 2017). Lastly, four repeated-measures Pearson correlation coefficients were calculated to test the strength of the relations between the mean normalized EMG amplitude and mean discharge rate at each contraction intensity for both decomposition approaches using the rmcorr package (Bakdash & Marusich, 2017). Data are presented as mean (standard deviation) below unless stated otherwise.

## RESULTS

### Trial exclusion

A total of 25 (1) [min-max: 24-29] submaximal voluntary contractions per participant were recorded and reduced to 17 (2) [min-max: 15-21] based on the exclusion criteria for motor unit decomposition. An average of 3 (1) trials [min-max: 2-5] were analysed per condition.

### Active torque

The maximum active dorsiflexion torque at the reference angles of -5, 20 and 45° plantar flexion were respectively 41.3 (9.7) Nm, 41.0 (9.6) Nm and 27.5 (6.9) Nm. As torque was an independent variable that was required to be matched by participants among conditions during the steady state, the effect of contraction intensity on active torque was significant (contraction intensity difference: 8.0 (1.8) Nm, *p*<.001), explaining 66.9% of the variance in active torque. There was no interaction between contraction intensity and condition for active torque and the effect of condition on active torque was also not significant (condition differences: ≤0.1 (0.2) Nm, *p*≥.054), explaining <0.2% of the variance in active torque. The magnitudes of the active torque difference among conditions at each contraction intensity were also not meaningful, falling within the measurement error of the torque sensor. The random effects estimates revealed that the active torque at 20% MVT in REF had the highest variance between participants (3.3 Nm^2^), followed by the active torque response to contraction intensity (3.1 Nm^2^). In contrast, condition explained much less of the variance in active torque (<0.1 Nm^2^) and unsurprisingly, participants who had higher active torques at 20% MVT also had higher active torques at 40% MVT (*r*=.99).

### Torque steadiness

Against our expectations, the effects of contraction intensity and condition on torque steadiness were not significant (contraction intensity difference: 0.14 (0.60)%, *p*=.342; condition differences: 0.08 (0.20)%, *p*≥.090), explaining only 1.5 and <0.2% of the variance in torque steadiness, respectively. There was no interaction between contraction intensity and condition for torque steadiness [min-max: 1.56-1.81%]. The random effects estimates revealed that contraction intensity was the largest source of variance in torque steadiness between participants (0.3%^2^) and surprisingly, participants who had higher torque steadiness at 20% MVT generally had lower torque steadiness at 40% MVT (*r*=-.83).

### Normalized EMG amplitude

The effects of contraction intensity and condition on normalized TA EMG amplitude were significant (*p*<.001 and *p*=.001, respectively), explaining 37.1 and 3.7% of the variance in EMG amplitude, respectively. There was also an interaction between contraction intensity and condition (*p*=.014), explaining 18.5% of the variance in EMG amplitude. As contraction intensity increased from 20 to 40% MVT, EMG amplitude increased (range of mean differences: 11.4 (2.4)% MVC, *t*(5015)=19.48, *p*<.001 to 13.6 (3.5)% MVC, *t*(5015)=15.97, *p*<.001, Table 1 and Fig. 2). As expected, EMG amplitude was also lower in STR_hold_ vs. REF at both contraction intensities, and EMG amplitude was higher in SHO_hold_ vs. REF at both contraction intensities (Table 1 and Fig. 2). Additionally, as expected, the amplitude difference between contraction intensities for SHO_hold_ was higher relative to REF (difference: 2.0 (2.7)% MVC), but similar for STR_hold_ (difference: 0.2 (2.7)% MVC; Table 1 and Fig. 2).

**Figure 2.**
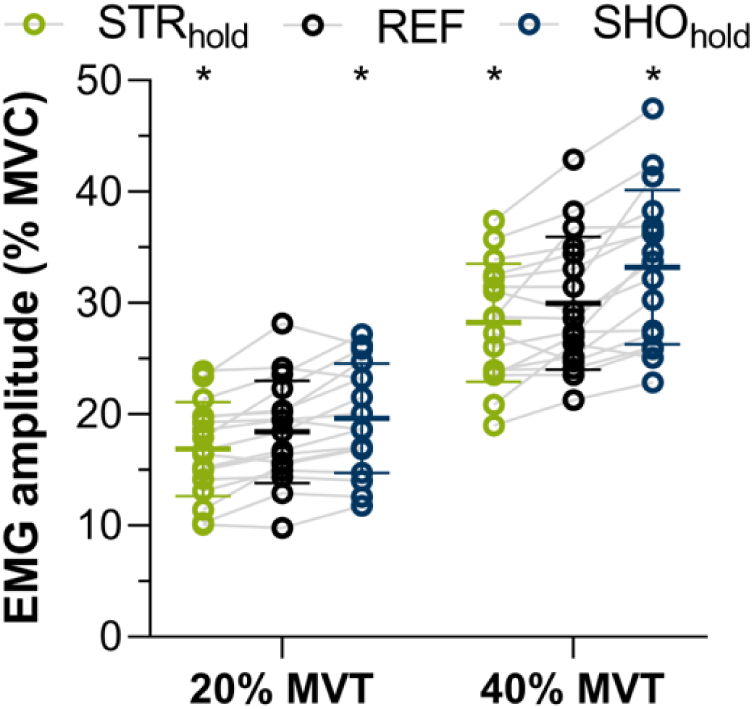
Normalized tibialis anterior EMG amplitude at 20 and 40% of maximum voluntary torque (MVT) during the steady-state phase in three conditions: stretch-hold (STR_hold_) contractions, fixed-end reference (REF) contractions, and shortening-hold contractions (SHO_hold_). Individual means (dots), between-group means (thick horizontal lines), and standard deviations (thin horizontal lines) are shown, with grey lines linking the conditions of each participant. Asterisks indicate a significant mean difference between the respective condition and REF (*p*≤.008).

**Table 1.** Normalized tibialis anterior EMG amplitudes during the torque-matched steady state of the six tested conditions.

|  | 20% MVT |  |  |  | 40% MVT |  |  |  |
| --- | --- | --- | --- | --- | --- | --- | --- | --- |
|  | EMG (% MVC) | Paired difference | *t* | *p* | EMG (% MVC) | Paired difference | *t* | *p* |
| Stretch-hold | 16.8 (4.2) | -1.6 (1.6)* | -3.88 | <.001 | 28.2 (5.3) | -1.7 (2.7)* | -2.64 | .008 |
| Reference | 18.4 (4.6) |  |  |  | 30.0 (6.0) |  |  |  |
| Shortening-hold | 19.6 (4.9) | 1.2 (1.9)* | 2.64 | .008 | 33.2 (6.9) | 3.2 (2.7)* | 4.87 | <.001 |

Values are between-subject means (standard deviations); $n=17$ . MVT, maximum voluntary torque; MVC, maximal voluntary contraction. \*Different to reference condition based on a linear mixed-effects model with Holm-adjustment for 3 tests.

### Decomposition output

The number of identified motor units and discharges differed between decomposition algorithms and contraction intensities (Table 2), as did the accuracy of the decomposition output (Table 3). However, at the same contraction intensity, both decomposition outputs shared most of their motor units [min-max: 67-100%] (Table 2), as expected due to the similar decomposition approaches. Also, unsurprisingly, the number of shared motor units between contraction intensities was low, at 18 (8)% [min-max: 0-29%] and 18 (11)% [min-max: 0-42%] for DEMUSE and MUedit, respectively; justifying the use of separate decompositions at each contraction intensity.

**Table 2.** Number of motor units and identified discharges between participants for each algorithm and shared between algorithms at each contraction intensity and combined. The shared row under discharge # indicates the % of shared motor units.

|  | Combined |  | 20% MVT |  | 40% MVT |  |
| --- | --- | --- | --- | --- | --- | --- |
|  | Total MU # | Total discharge # | MU # | Discharge # | MU # | Discharge # |
| DEMUSE | 503 | 340581 | 16 (5) [8-31] | 10103 (3958) | 13 (5) [5-22] | 9932 (3614) |
| MUedit | 777 | 512492 | 25 (9) [9-37] | 14714 (6135) | 21 (9) [9-34] | 15432 (6631) |
| Shared | 456 | 91% | 15 (6) [6-30] | 91 (9) [67-100%] | 12 (5) [4-21] | 87 (10) [67-100%] |

Values are between-subject means (standard deviations) [minimum-maximum] except for combined, which is the between-subject sum; $n=17$ . MU, motor unit; MVT, maximum voluntary torque.

**Table 3.** Pulse-to-noise ratios and silhouette values of the identified motor units between participants for each algorithm at each contraction intensity.

|  | 20% MVT |  |  | 40% MVT |  |  |
| --- | --- | --- | --- | --- | --- | --- |
|  | PNR (dB) | SIL | CV in DR (%) | PNR (dB) | SIL | CV in DR (%) |
| DEMUSE | 32 (3) [28-37] | .93 (.02) [.87-.96] | 13 (2) [10-18] | 31 (2) [27-35] | .92 (.02) [.88-.94] | 15 (3) [10-21] |
| MUedit | 34 (2) [31-38] | .93 (.01) [.91-.95] | 18 (2) [15-22] | 33 (2) [30-35] | .92 (.01) [.91-.94] | 20 (2) [16-26] |

Values are between-subject means (standard deviations) [minimum-maximum]; $n=17$ . MVT, maximum voluntary torque; PNR, pulse-to-noise ratio; SIL, silhouette value, CV, coefficient of variation; DR, discharge rate.

### Discharge rate

The effect of the decomposition algorithm on tibialis anterior motor unit discharge rate was not significant (decomposition difference: 0.01 (0.33) Hz, *p*=.871), explaining 0% of the variance in motor unit discharge rate. Consequently, the estimated marginal means reported below were averaged between decompositions. The effects of contraction intensity and condition on discharge rate were significant (both *p*<.001), explaining 25.1 and 7.5% of the variance in discharge rate, respectively. There was also a significant interaction between contraction intensity and condition (*p*=.002) that explained 14.8% of the variance in discharge rate. As contraction intensity increased from 20 to 40% MVT, discharge rate increased (range of mean differences: 2.3 (0.9) Hz, *t*(17.4)=10.95, *p*<.001 to 3.3 (0.8) Hz, *t*(14.0)=16.07, *p*<.001; Table 4 and Fig. 3). As expected, discharge rate was also lower in STR_hold_ vs. REF at both contraction intensities, but surprisingly, discharge rate was only higher in SHO_hold_ vs. REF at 40 and not 20% MVT (Table 4 and Fig. 3).

**Figure 3.**
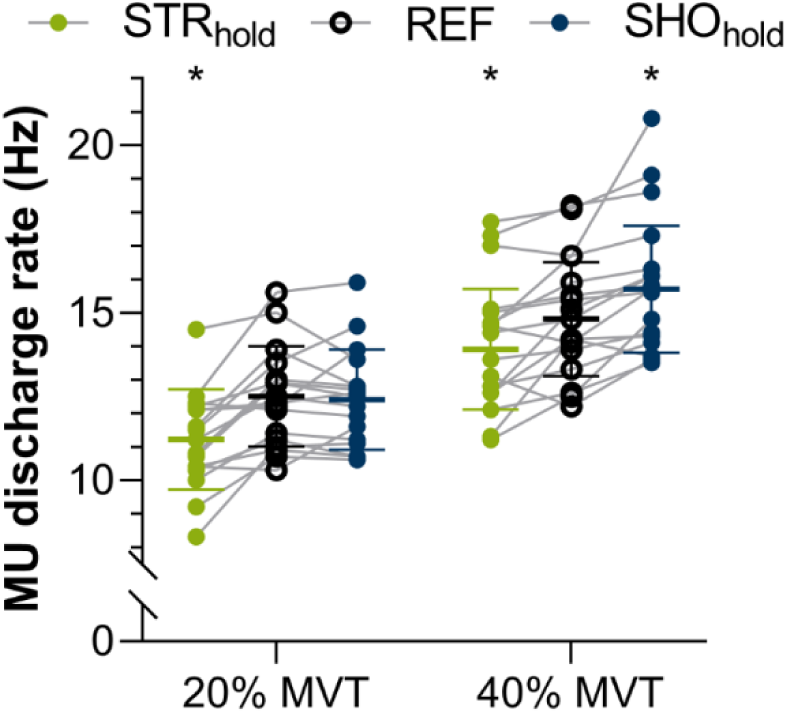
Tibialis anterior motor unit (MU) discharge rates at 20 and 40% of maximum voluntary torque (MVT) during the steady-state phase in three conditions: stretch-hold (STR_hold_) contractions, fixed-end reference (REF) contractions, and shortening-hold contractions (SHO_hold_). Individual means (dots), between-group estimated marginal means (thick horizontal lines) and associated standard deviations (thin horizontal lines) are shown, with grey lines linking the conditions of each participant. Asterisks indicate a significant mean difference between the respective condition and REF (*p*≤.004).

**Table 4.** Tibialis anterior motor unit discharge rates averaged from the two decomposition algorithms (DEMUSE and MUedit) during the torque-matched steady state of the six conditions.

|  | 20% MVT |  |  |  | 40% MVT |  |  |  |
| --- | --- | --- | --- | --- | --- | --- | --- | --- |
|  | Discharge rate (Hz) | Paired difference | *t* | *p* | Discharge rate (Hz) | Paired difference | *t* | *p* |
| Stretch-hold | 11.2 (1.5) | -1.3 (1.0)* | -5.07 | <.001 | 13.9 (1.8) | -0.8 (1.0)* | -3.35 | .004 |
| Reference | 12.5 (1.5) |  |  |  | 14.8 (1.7) |  |  |  |
| Shortening-hold | 12.4 (1.5) | 0.0 (0.8) | -0.04 | .966 | 15.7 (1.9) | 1.0 (1.0)* | 3.85 | .003 |

Values are between-subject means (discharge rate column) and within-subject mean differences (paired difference column) (standard deviations); *n*=17. MVT, maximum voluntary torque. \*Different to reference condition based on a linear mixed-effects model with Holm-adjustment for 3 tests.

The random effects estimates revealed that contraction intensity was the largest source of variance at the motor unit level (3.2 Hz^2^), followed by decomposition algorithm at the motor unit level (2.9 Hz^2^). Consequently, the discharge rate response to contraction intensity and decomposition algorithm was not consistent between motor units within the same participant and this effect was larger at 40 than 20% MVT and for MUedit than DEMUSE (1.5 Hz^2^). Less of the variance in discharge rate was explained by contraction intensity and decomposition algorithm at the participant level (0.5 and <0.1 Hz^2^, respectively), indicating that the effect of both factors on discharge rate was more consistent when averaged between motor units within each participant than at the motor unit level. The opposite was true for condition, whereby more of the variance was explained by condition at the participant level (STR_hold_ vs. SHO_hold_: 0.9 vs. 0.5 Hz^2^) than at the motor unit level (<0.1 Hz^2^). Consequently, the discharge rate response to condition was more consistent at the motor unit level than when averaged between motor units within each participant.

The repeated-measures linear relations between normalized TA EMG amplitude (see Fig. 4 for representative time traces) and TA discharge rate were strong and positive at each contraction intensity and with each decomposition approach; albeit the relations were slightly weaker at 20 (DEMUSE: *r*_rm_(33)=.84 [.70 to .92], *p*<.001; MUedit: *r*_rm_(33)=.85 [.72 to .92], *p*<.001) vs. 40% MVT (DEMUSE: *r*_rm_(33)=.92 [.85 to .96], *p*<.001; MUedit: *r*_rm_(33)=.94 [.88 to .97], *p*<.001) because discharge rate did not increase in line with EMG amplitude at 20% MVT in SHO_hold_.

**Figure 4.**
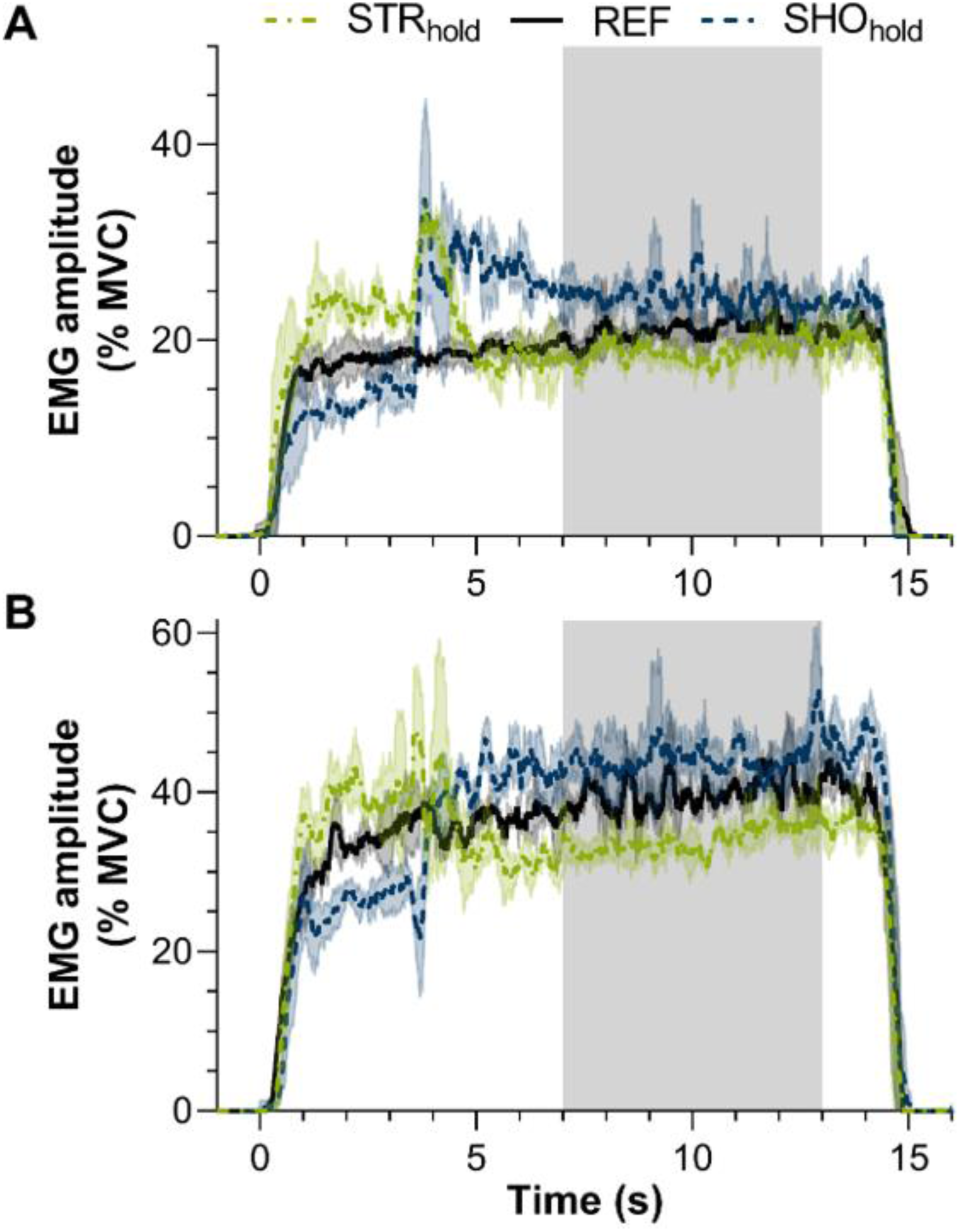
Normalized tibialis anterior EMG amplitudes over time at (A) 20 and (B) 40% of maximum voluntary torque in three conditions: stretch-hold (STR_hold_) contractions, fixed-end reference (REF) contractions, and shortening-hold contractions (SHO_hold_). The mean trace of multiple trials per condition from one participant (same as in Fig. 1) are shown, with shaded areas around the mean indicating the standard deviation and the grey shaded area indicating the 6-s steady-state phase. The normalized EMG amplitude was calculated as the mean of the 59 single-differential EMG signals previously normalized to the channel-specific maximal EMG amplitude from the MVC with the highest active torque at 20° plantar flexion.

### Discharge rate variability

The effect of decomposition on motor unit discharge rate variability was significant (decomposition difference: 4.38 (1.29)%, *p*<.001, Table 3), explaining 14.1% of the variance in discharge rate variability, with MUedit having a higher CV than DEMUSE. There was no interaction between contraction intensity and condition for discharge rate variability [min-max: 14.9-17.9%]. The effect of contraction intensity on discharge rate variability was significant (*p*=.004), explaining 2.8% of the variance in discharge rate variability; with increased variability at 40 vs. 20% MVT (1.86 (2.28)%, Table 3). However, against our expectations, the effect of condition on discharge rate variability was not significant (p=.249), explaining only 0.9% of the variance in discharge rate variability.

### Unmatched motor units

By design, the number of matched motor units analysed was consistent among conditions at each contraction intensity. However, pre-analysis, there were more instances when additional (i.e. non-matched) motor units were identified in SHO_hold_ than in REF and STR_hold_ at both contraction intensities, independent of the decomposition algorithm (Fig. 5). Additionally, there were more instances when additional motor units were identified in REF than STR_hold_ (Fig. 5), but the variability between participants was high (DEMUSE: 5 and 3 from 17 participants had additional motor units identified in SHO_hold_ and REF at 20 and 40% MVT, respectively; MUedit: 10 and 6 respective participants exhibited this pattern).

**Figure 5.**
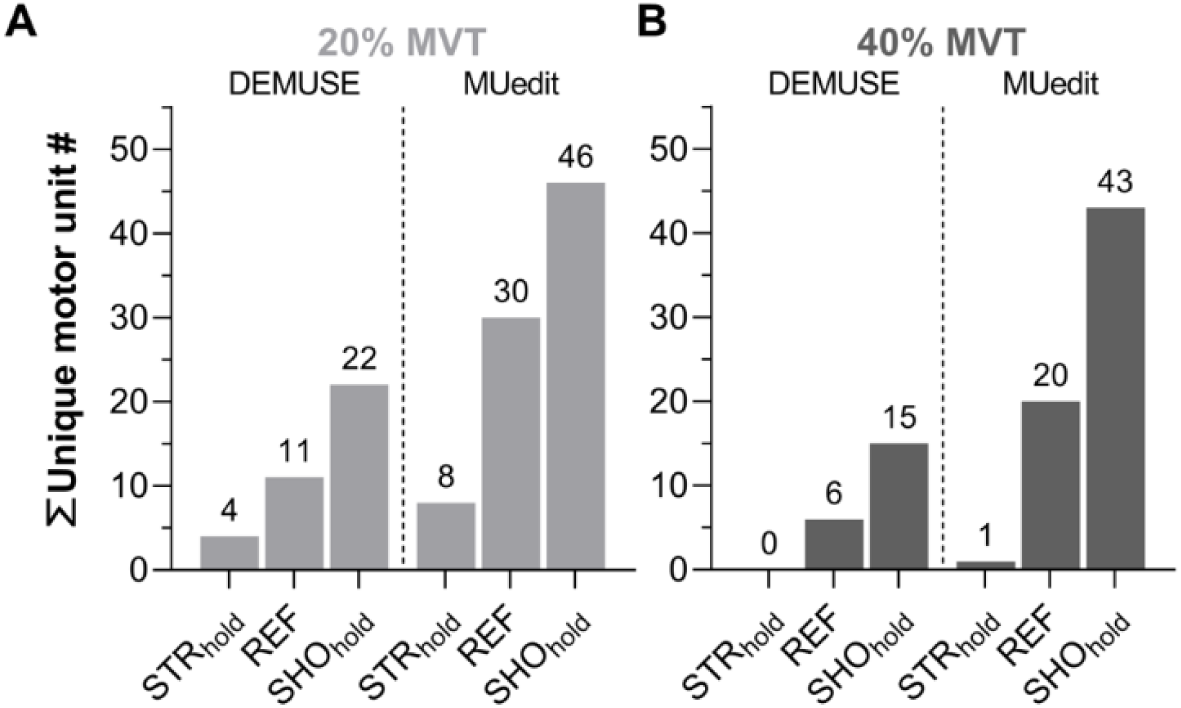
Between-participant sum of unique (i.e. non-matched) tibialis anterior motor units with each decomposition approach (DEMUSE or MUedit) at (A) 20 or (B) 40% of maximum voluntary torque (MVT) during the steady-state phase in three conditions: stretch-hold (STR_hold_) contractions, fixed-end reference (REF) contractions, and shortening-hold contractions (SHO_hold_).

## DISCUSSION

The present study was designed to determine the extent of neural drive modulation within the human TA induced following active muscle stretch or shortening compared with torque- and joint-angle-matched fixed-end reference contractions. As expected, we found that neural drive decreased similar amounts following active stretch at two contraction intensities (20 and 40% MVT), whereas neural drive increased more following active shortening at 40% MVT, due to increases in discharge rate that were not apparent at 20% MVT. These results suggest that the neural drive modulation following active shortening, but not active stretch, is contraction-intensity specific, presumably to account for differences in rFD between contraction intensities (De Ruiter *et al*., 1998; Raiteri *et al*., 2024) that might not exist for rFE (Oskouei & Herzog, 2006; Seiberl *et al*., 2012). We also found indirect evidence of additional motor unit recruitment following active shortening and reduced motor unit recruitment following active stretch based on a respectively higher and lower number of unique motor units relative to the reference condition at both contraction intensities. Against our expectations, we did not find increased torque variability following active stretch contractions, which might be due to the shorter muscle-tendon unit length we tested at and correspondingly less rFE (Bakenecker *et al*., 2022) compared with previous work (Jakobi *et al*., 2020). In combination, our findings suggest that EMG-amplitude-based muscle force predictions will likely be inaccurate following an active muscle length change due to changes in the muscle’s neural drive to account for changes in its force capacity.

### Neural drive modulations

Following active stretch, we observed fewer unique motor units at 20 and 40% MVT, and similar respective decreases of 9 and 6% in EMG amplitude and 10 and 6% in discharge rate of matched motor units relative to the reference condition. Such reductions in EMG amplitude fall within the range of 5 to 44% from previous in vivo torque-matched experiments on the TA (Paquin & Power, 2018; Jakobi *et al*., 2020), soleus (Dalton *et al*., 2018), superficial quadriceps (Altenburg *et al*., 2008) and adductor pollicis (Oskouei & Herzog, 2005). Like the current study, these studies (except one; Altenburg *et al*., 2008) permitted increases in torque during active stretch. If we therefore ignore Altenburg et al. (2008) and compare the other studies with contraction intensities from 20 to 40% MVT, reductions in EMG amplitude ranged from 5 (Dalton *et al*., 2018) and 12% (Oskouei & Herzog, 2005) to 24 (Paquin & Power, 2018; Jakobi *et al*., 2020) and 35% (Paquin & Power, 2018). Consequently, the previous studies on TA indicate larger reductions in EMG amplitude following active stretch compared with the current study, which might be because: 1) we did not exclude “non-responders” from the statistical analysis as all participants exhibited EMG amplitude reductions following stretch in some trials, and; 2) we also tested at a comparably shorter reference muscle-tendon unit length. As rFE appears to increase with an increase in the final muscle length attained during submaximal voluntary contractions (Bakenecker *et al*., 2022), it is possible that our shorter reference length (i.e. 20° plantar flexion versus 25 or 40° plantar flexion) led to less rFE than previous studies on the TA (Paquin & Power, 2018; Jakobi *et al*., 2020), and correspondingly attenuated the reduction in EMG amplitude following active stretch. Notably, we did not test at a relatively longer reference length though because we wanted to match the ranges of motion of the stretch and shortening contractions.

Our observed reduction in discharge rate of 10% following active stretch at 20% MVT is likely less than the 27% reduction reported by Jakobi and colleagues (2020) at 20% MVT for the reasons outlined above. However, our discharge rate findings are also at odds with vastus lateralis findings reported by Altenburg and colleagues (2008), who found non-significant reductions in discharge rate of 1% following active stretch. Again, this might be because these authors (Altenburg *et al*., 2008) tested at a comparably shorter reference length (i.e. on the plateau versus descending limb of the muscle’s force-length relation), which may have lead to less rFE following active stretch. But the torque-matching strategy by Altenburg and colleagues (2008) was also different, as participants were instructed to match a constant torque during active muscle-tendon unit stretch. This torque-matching strategy may have reduced the rFE following active stretch, because rFE and peak fascicle force during stretch are positively correlated; at least within the TA during maximal voluntary contractions (95% CI: .30 to .82; Tecchio *et al*., 2024). Consequently, the amount of neural drive modulation following active stretch probably depends on the peak force attained during stretch and the reference muscle length.

Following active shortening, we observed additional unique motor units at 20 and 40% MVT, and dissimilar respective increases of 7 and 11% in EMG amplitude and 0 and 7% in discharge rate of matched motor units relative to the reference condition. Such increases in EMG amplitude fall within the range of 9 to 48% from previous in vivo torque-matched experiments on the TA (Paquin & Power, 2018), superficial quadriceps (Altenburg *et al*., 2008) and adductor pollicis (Rousanoglou *et al*., 2007). However, the larger increase in neural drive at 40 versus 20% MVT in the current study is at odds with one previous study, which found similar increases in EMG amplitude (9-25%) and discharge rate (3-10%) between contraction intensities ranging from 4 to 50% MVT (Altenburg *et al*., 2008). This difference might again arise because of differences in torque-matching strategies as explained above, as well as reduced statistical power in the previous study due to an unclear systematic difference between contraction intensities (4-47% MVT) and a reduced mean motor unit yield of ∼4 per person from intramuscular EMG (Altenburg *et al*., 2008).

More recent findings from Paquin and Power (2018) on the TA support those of our study, as these authors reported increasing rFD effect sizes with increasing contraction intensity. However, Paquin and Power (2018) used conventional bipolar surface EMG and thus could not provide data-driven insights into the neural drive modulation strategies following active shortening. Interestingly, we found that the neural drive modulation strategy differed between contraction intensities, with discharge rate only increasing at 40% MVT. The lack of discharge rate modulation at 20% MVT and corresponding increase in EMG amplitude following active shortening, as well as the additional unique motor units detected compared with the reference condition, indicate that additional motor unit recruitment was the primary strategy employed by the central nervous system to overcome the rFD following shortening. A motor unit recruitment strategy may be ideal as it recruits new muscle fibres that do not have depressed force capacities due to active shortening, and this strategy also prevents additional active cycling of depressed cross bridges within the already-recruited fibres. The neural drive modulation strategy at 20% MVT was clearly different to the combined motor unit recruitment and discharge rate modulation strategy observed following shortening at 40% MVT, where it is unclear how much the additional increase in discharge rate helped to compensate for additional rFD; with additional rFD being likely following higher-force shortening based on previous findings from the cat soleus (Leonard & Herzog, 2005; Herzog & Leonard, 2007), human adductor pollicis (De Ruiter & De Haan, 2003) and human TA (Raiteri *et al*., 2024). A combined motor unit recruitment and discharge rate modulation strategy to account for rFD at higher contraction intensities might not be preferable, but rather naturally manifest due to the increased common synaptic input to alpha motoneurons, which limits the independent control of individual motor units within the active pool (Farina *et al*., 2014), leading to an inevitable increase in discharge rate. Consequently, we speculate that there might be unexplored constraints on the strategies used by the central nervous system to compensate for rFD following shortening that are dependent on the strength of neural drive and rFD magnitude.

In line with our third hypothesis, we found that repeated-measures changes in TA motor unit discharge rates and EMG amplitudes were strongly and positively correlated at 20 and 40% MVT (95% CI with decompositions averaged: .71-.92 and .88-.97, respectively). However, the correlation was not perfect, which is to be expected because EMG amplitudes should increase with increases in discharge rate of the same motor units across conditions, as well as with additional motor unit recruitment (Merletti & Farina, 2016). Due to the low and variable number of additional (i.e. unique) motor units detected (Fig. 5), we did not statistically analyse the discharge rates of unique motor units; but based on random chance we would expect a 33% chance of detecting unique motor units in each condition. However, we found respective distributions in STR_hold_, REF and SHO_hold_ of 10, 34 and 56% at 20% MVT, and 1, 31, and 68% at 40% MVT, which suggests additional motor unit recruitment in SHO_hold_ relative to REF, and reduced motor unit recruitment in STR_hold_ relative to REF. Although this pattern varied between individuals, this observation indicates that comparing discharge rates of matched motor units only leads to a loss of information about unique motor units and potential differences in motor unit recruitment among conditions. Indeed, this loss of information is reflected by worse predictions of contraction-intensity-dependent changes in torque with discharge rates from matched motor units (.84-.98) than EMG amplitudes (.91-.99).

### Torque steadiness

We did not find support for our hypothesis that torque steadiness was reduced following muscle-tendon unit stretch compared with following shortening or during the fixed-end reference contractions. As discussed above, this could be because we induced less rFE at our relatively shorter reference TA muscle-tendon unit length compared with the study by Jakobi and colleagues (2020). Additionally, the discrepancy between our results might be explained by recent findings from our group; we observed that large torque differences during a contraction can reduce torque steadiness (Raiteri et al. 2026 [unpublished]), and in the previous study (Jakobi *et al*., 2020) the torque difference was ∼4-fold at the start versus end of the STR_hold_ contractions, unlike in our study (Fig. 1A). However, it is also worth noting that the previously observed mean reduction in torque steadiness following stretch was smaller (1% with an unreported standard deviation) at 20 than 10% MVT (3%), and torque steadiness has been reported to decrease from 20 to 5-10% MVT, at least during isometric finger abduction (Galganski *et al*., 1993) and trunk flexion (Deering *et al*., 2017). The previously reported ∼3-fold difference in the torque steadiness reduction (Jakobi *et al*., 2020) is thus not well explained by rFE-based mechanisms as rFE appears to be independent of contraction intensity (Oskouei & Herzog, 2006; Seiberl et al., 2012). Therefore, additional studies at different contraction intensities and final MTU lengths are needed to confirm whether rFE-based mechanisms drive reductions in torque steadiness following active muscle stretch.

### Sex differences

We did not use null-hypothesis testing to investigate sex differences in the current study because only a very large effect size of greater than 1.96 (Cohen’s *d*_z_) could have been detected with a two-tailed alpha level of 5% and a desired statistical power of 95% with six women and eleven men. With that in mind, there appeared to be 3 to 7 fewer motor units identified per person in women compared with men, which is in line with some previous reports (Taylor *et al*., 2022; Lulic-Kuryllo & Inglis, 2022; Jenz *et al*., 2023), but not others (Inglis & Gabriel, 2020, 2021; Yacyshyn *et al*., 2025; Luckey *et al*., 2026). We also observed greater sex differences in the motor unit yield with MUedit than DEMUSE (Table 5), which might help to explain discrepancies between previous studies. The mean tibialis anterior motor unit discharge rates and discharge rate variability were also slightly higher in women than men (0.8-0.9 Hz and 0.5%; Table 6), which is similar to previous reports of higher TA motor unit discharge rates (Kowalski & Christie, 2020; Inglis & Gabriel, 2020; Taylor *et al*., 2022) and variability (Kowalski & Christie, 2020; Inglis & Gabriel, 2021) in women than men at similar contraction intensities. However, it is unclear if these differences in our study are meaningful, as we also found higher normalized EMG amplitudes in women than men (1.8% MVC; Table 6), which might have led to the slightly higher discharge rates and variability in women. Clearly, future studies with at least 27-105 women and the same number of men are needed to confirm whether there are modest to large (Cohen’s *d*_z_ ≥0.5-≥1) sex differences in motor unit behaviour.

**Table 5.** Number of motor units identified between sexes for each algorithm at each contraction intensity and combined.

|  | Combined<br>Total MU # |  | 20% MVT<br>MU # |  | 40% MVT<br>MU # |  |
| --- | --- | --- | --- | --- | --- | --- |
|  | Women | Men | Women | Men | Women | Men |
| DEMUSE | 148 | 355 | 14 (5) [8-20] | 18 (5) [12-31] | 11 (5) [6-19] | 14 (4) [5-22] |
| MUedit | 217 | 560 | 20 (11) [9-35] | 27 (7) [17-37] | 17 (9) [9-34] | 24 (8) [9-33] |

Values are between-subject means (standard deviations) [minimum-maximum] except for combined, which is the between-subject sum; $n=6$ (women) and $n=11$ (men). MU, motor unit; MVT, maximum voluntary torque.

**Table 6.** Sex differences in tibialis anterior motor unit discharge rates, discharge rate variability and normalized tibialis anterior EMG amplitudes during the torque-matched steady state of the six tested conditions. Discharge rate and discharge rate variability were averaged from the two decomposition algorithms (DEMUSE and MUedit), and all three variables were averaged among conditions.

|  | Discharge rate<br>(Hz) |  | Discharge rate<br>variability (%) |  | EMG (% MVC) |  |
| --- | --- | --- | --- | --- | --- | --- |
|  | 20% MVT | 40% MVT | 20% MVT | 40% MVT | 20% MVT | 40% MVT |
| Women | 12.6 (1.9) | 15.4 (2.0) | 15.8 (2.2) | 17.9 (2.6) | 19.5 (5.9) | 31.6 (6.6) |
| Men | 11.8 (1.2) | 14.5 (1.3) | 15.3 (1.7) | 17.4 (2.1) | 17.7 (4.3) | 29.8 (5.0) |

Values are between-subject means (standard deviations); $n=6$ (women) and $n=11$ (men). MVT, maximum voluntary torque; MVC, maximal voluntary contraction.

### Limitations

The neural drive changes we observed are likely muscle-length specific, since rFE and rFD are both affected by muscle length (Van Noten & Van Leemputte, 2011; Bakenecker *et al*., 2022). Due to our study design, we were limited to comparing motor unit discharge rates because each condition started at a different muscle length, preventing us from attaining other important insights, such as differences in motor unit recruitment thresholds. We were also limited to decomposing only the steady state of the contractions due to current limitations with tracking temporal and spatial changes in the waveforms of the surface-recorded motor unit action potentials. Further, it is important to note that contraction intensity differences during the active stretch of shortening phases of our experiments might have influenced the motor unit firing properties during the following steady-state phase, independent of changes due to active stretch or shortening (i.e. if dynamic torques were matched, the neural drive reduction or increase may be have attenuated or intensified following stretch or shortening, respectively). Lastly, we limited the number of decomposition iterations to 50 to limit processing time to around 10 minutes because 250 iterations with MUedit did not just increase the processing time by a factor of 5, but by a factor of 26 to 54 (i.e. 4 to 9 hours), without a notable increase in the motor unit yield following automatic editing.

Our finding that the discharge rate response to contraction intensity and decomposition algorithm was more variable at the motor unit level than at the participant level (i.e. motor units averaged within each participant), while the reverse was true for condition, is likely because we performed separate decompositions at each contraction intensity. We found that grouping the contraction intensities together biased the decomposition to higher-threshold motor units active at 40 but not 20% MVT, subsequently reducing the motor unit yield at 20% MVT, which is likely because decompositions are biased to identify sources with the highest energy (Negro *et al*., 2016). Our approach to overcome this limitation, by performing contraction-intensity-specific decompositions, likely explains why motor units less consistently responded to contraction intensity than condition, i.e. we tracked different motor units between contraction intensities, but the same motor units among conditions. Consequently, the opposite effects of contraction intensity and condition on the discharge rate response at the motor unit level versus motor-unit-averaged participant level are not surprising.

### Relevance

Our results add to the growing body of evidence that muscle force predictions based on EMG measurements are likely to be inaccurate following dynamic movements because active muscle stretch and shortening lead to rFE and rFD, which alter the neural drive to a muscle to attain a given net joint torque. Although the direction (i.e. overestimation or underestimation) of force error is presumably easy to predict based on the direction of muscle length change, the magnitude of this error is, based on our data, not consistent between contraction intensities or participants, which indicates that a contraction-intensity-specific and subject-specific calibration is required to improve EMG-based muscle force predictions. Further work should investigate other muscles and the effect of different preloads, ranges of motion and speeds, as well as whether neural drive changes during active stretch or shortening mirror or oppose those following active stretch or shortening.

## Conclusions

In conclusion, we found that to account for the enhanced force capacity of muscle following active stretch, neural drive to the TA is reduced to attain a given dorsiflexion torque. Conversely, to account for the depressed force capacity of muscle following active shortening, neural drive to the TA is increased. These findings are based on a robust two-method decomposition approach yielding 17 to 21 decomposed motor units per person on average and suggest that the neural drive to a muscle depends on the muscle’s history-dependent force capacity. Importantly, the way neural drive is modulated differs among contraction types, being independent of contraction intensity following active muscle stretch, but not following active muscle shortening. Inherently, the neural drive strategy used by the central nervous system to account for force capacity changes following active shortening appears ideal at a low contraction intensity as force is increased via the recruitment of new muscle fibres without depressed force capacities, whereas at a relatively higher contraction intensity, the strategy is less preferable. A combined motor unit recruitment and discharge rate modulation strategy to account for additional rFD following higher shortening forces might be due to constraints imposed by increasing common synaptic input to the alpha motoneurons, which might make overcoming force deficits following active shortening contractions more problematic for individuals with strength deficits.

## ACKNOWLEDGEMENTS

This study was supported by the DFG (447345165). We thank Dr. Alberto Botter (Turin, Italy) for supervising the manual editing of the DEMUSE decompositions and for his insightful suggestions. We also thank Dr. Simon Avrillon (Nantes, France) and Marius Oßwald (Erlangen, Germany) for contributing MATLAB code for the decomposition comparisons.

## AUTHOR CONTRIBUTIONS

B.J.R., K.F.B. and D.H. conceived and designed research, K.F.B. performed experiments, B.J.R. analyzed data, M.B. manually edited DEMUSE decompositions, A.C.V. assisted with the statistical analysis and interpretation, B.J.R. prepared figures, B.J.R. drafted manuscript, all authors interpreted results of experiments, all authors edited and revised manuscript, all authors approved final version of manuscript.

## DATA AVAILABILITY

Datasets generated and analyzed during the current study are available in the following Zenodo repository [https://doi.org/10.5281/zenodo.18922801].

## REFERENCES

Abbott BC & Aubert XM (1952). The force exerted by active striated muscle during and after change of length. J Physiol 117, 77–86.

Altenburg TM, De Ruiter CJ, Verdijk PWL, Van Mechelen W & De Haan A (2008). Vastus lateralis surface and single motor unit EMG following submaximal shortening and lengthening contractions. Appl Physiol Nutr Metab 33, 1086–1095.

Avrillon S, Hug F, Baker SN, Gibbs C & Farina D (2024). Tutorial on MUedit: An open-source software for identifying and analysing the discharge timing of motor units from electromyographic signals. J Electromyogr Kinesiol 77, 102886.

Bakdash JZ & Marusich LR (2017). Repeated measures correlation. Front Psychol 8, 1–13.

Bakenecker P, Weingarten T, Hahn D & Raiteri BJ (2022). Residual force enhancement is affected more by quadriceps muscle length than stretch amplitude. Elife 11, e77553.

Bates D, Mächler M, Bolker B & Walker S (2015). Fitting linear mixed-effects models using **lme4**. J Stat Softw; DOI: 10.18637/jss.v067.i01.

Brand RA, Pedersen DR & Friederich JA (1986). The sensitivity of muscle force predictions to changes in physiologic cross-sectional area. J Biomech 19, 589–596.

Caillet AH, Phillips ATM, Modenese L & Farina D (2024). NeuroMechanics: Electrophysiological and computational methods to accurately estimate the neural drive to muscles in humans in vivo. J Electromyogr Kinesiol 76, 102873.

Chen M & Zhou P (2016). A novel framework based on fastICA for high density surface EMG decomposition. IEEE Trans Neural Syst Rehabil Eng 24, 117–127.

Dalton BH, Contento VS & Power GA (2018). Residual force enhancement during submaximal and maximal effort contractions of the plantar flexors across knee angle. J Biomech 78, 70–76.

De Ruiter CJ & De Haan A (2003). Shortening-induced depression of voluntary force in unfatigued and fatigued human adductor pollicis muscle. J Appl Physiol 94, 69–74.

De Ruiter CJ, De Haan A, Jones DA & Sargeant AJ (1998). Shortening-induced force depression in human adductor pollicis muscle. J Physiol 507, 583–591.

Deering RE, Senefeld JW, Pashibin T, Neumann DA & Hunter SK (2017). Muscle function and fatigability of trunk flexors in males and females. Biol Sex Differ 8, 12.

Del Vecchio A, Holobar A, Falla D, Felici F, Enoka RM & Farina D (2020). Tutorial: Analysis of motor unit discharge characteristics from high-density surface EMG signals. J Electromyogr Kinesiol 53, 102426.

Dideriksen JL & Farina D (2019). Amplitude cancellation influences the association between frequency components in the neural drive to muscle and the rectified EMG signal ed. Haith AM. PLoS Comput Biol 15, e1006985.

Dimitrov GV, Arabadzhiev TI, Hogrel J-Y & Dimitrova NA (2008). Simulation analysis of interference EMG during fatiguing voluntary contractions. Part II – Changes in amplitude and spectral characteristics. J Electromyogr Kinesiol 18, 35–43.

Edman KA, Elzinga G & Noble MI (1978). Enhancement of mechanical performance by stretch during tetanic contractions of vertebrate skeletal muscle fibres. J Physiol 281, 139–155.

Farina D, Negro F & Dideriksen JL (2014). The effective neural drive to muscles is the common synaptic input to motor neurons. J Physiol 592, 3427–3441.

Faul F, Erdfelder E, Buchner A & Lang A-G (2009). Statistical power analyses using G*Power 3.1: Tests for correlation and regression analyses. Behav Res Methods 41, 1149–1160.

Fife D (2022). Flexplot: graphically-based data analysis. Psychol Methods 27, 477–496.

Galganski ME, Fuglevand AJ & Enoka RM (1993). Reduced control of motor output in a human hand muscle of elderly subjects during submaximal contractions. J Neurophysiol 69, 2108–2115.

Granzier HL & Pollack GH (1989). Effect of active pre-shortening on isometric and isotonic performance of single frog muscle fibres. J Physiol 415, 299–327.

Grison A, Mendez Guerra I, Clarke AK, Muceli S, Ibáñez J & Farina D (2025). Unlocking the full potential of high-density surface EMG: novel non-invasive high-yield motor unit decomposition. J Physiol 603, 2281–2300.

Herzog W & Leonard TR (2007). Residual force depression is not abolished following a quick shortening step. Journal of Biomechanics 40, 2806–2810.

Herzog W, Leonard TR & Wu JZ (1998). Force depression following skeletal muscle shortening is long lasting. J Biomech 31, 1163–1168.

Holobar A, Farina D, Gazzoni M, Merletti R & Zazula D (2009). Estimating motor unit discharge patterns from high-density surface electromyogram. Clin Neurophysiol 120, 551–562.

Holobar A, Minetto MA, Botter A, Negro F & Farina D (2010). Experimental analysis of accuracy in the identification of motor unit spike trains from high-density surface EMG. IEEE Trans Neural Syst Rehabil Eng 18, 221–229.

Holobar A, Minetto MA & Farina D (2014). Accurate identification of motor unit discharge patterns from high-density surface EMG and validation with a novel signal-based performance metric. J Neural Eng 11, 1–11.

Holobar A & Zazula D (2004). Correlation-based decomposition of surface electromyograms at low contraction forces. Med Biol Eng Comput 42, 487–495.

Holobar A & Zazula D (2007a). Multichannel blind source separation using convolution kernel compensation. IEEE Trans Signal Process 55, 4487–4496.

Holobar A & Zazula D (2007b). Gradient convolution kernel compensation applied to surface electromyograms. In Independent Component Analysis and Signal Separation, ed. Davies ME, James CJ, Abdallah SA & Plumbley MD, Lecture Notes in Computer Science, pp. 617–624. Springer Berlin Heidelberg, Berlin, Heidelberg. Available at: http://link.springer.com/10.1007/978-3-540-74494-8_77 [Accessed August 14, 2025].

Hug F, Avrillon S, Del Vecchio A, Casolo A, Ibanez J, Nuccio S, Rossato J, Holobar A & Farina D (2021). Analysis of motor unit spike trains estimated from high-density surface electromyography is highly reliable across operators. J Electromyogr Kinesiol 58, 102548.

Inglis JG & Gabriel DA (2020). Sex differences in motor unit discharge rates at maximal and submaximal levels of force output. Appl Physiol Nutr Metab 45, 1197–1207.

Inglis JG & Gabriel DA (2021). Sex differences in the modulation of the motor unit discharge rate leads to reduced force steadiness. Appl Physiol Nutr Metab 46, 1065–1072.

Jakobi JM, Kuzyk SL, McNeil CJ, Dalton BH & Power GA (2020). Motor unit contributions to activation reduction and torque steadiness following active lengthening: a study of residual torque enhancement. J Neurophysiol 123, 2209–2216.

Jenz ST, Beauchamp JA, Gomes MM, Negro F, Heckman CJ & Pearcey GEP (2023). Estimates of persistent inward currents in lower limb motoneurons are larger in females than in males. J Neurophysiol 129, 1322–1333.

Kalc M, Skarabot J, Divjak M, Urh F, Kramberger M, Vogrin M & Holobar A (2023). Identification of motor unit firings in H-reflex of soleus muscle recorded by high-density surface electromyography. IEEE Trans Neural Syst Rehabil Eng 31, 119–129.

Kowalski KL & Christie AD (2020). Force control and motor unit firing behavior following mental fatigue in young female and male adults. Front Integr Neurosci 14, 15.

Kuznetsova A, Brockhoff PB & Christensen RHB (2017). **lmerTest** package: tests in linear mixed effects models. J Stat Softw; DOI: 10.18637/jss.v082.i13.

Lenth RV, Piaskowski J, Banfai B, Bolker B, Buerkner P, Giné-Vázquez I, Hervé M, Jung M, Love J, Miguez F, Riebl H & Singmann H (2025). emmeans: Estimated Marginal Means, aka Least-Squares Means. Available at: https://cran.r-project.org/web/packages/emmeans/index.html [Accessed January 6, 2026].

Leonard TR, DuVall M & Herzog W (2010). Force enhancement following stretch in a single sarcomere. Am J Physiol Cell Physiol 299, C1398–C1401.

Leonard TR & Herzog W (2005). Does the speed of shortening affect steady-state force depression in cat soleus muscle? J Biomech 38, 2190–2197.

Lloyd DG & Besier TF (2003). An EMG-driven musculoskeletal model to estimate muscle forces and knee joint moments in vivo. J Biomech 36, 765–776.

Luckey CJ, Marsala MJ & Christie AD (2026). Sex-related differences in motor unit yield, subcutaneous tissue thickness, and maximal force. Appl Physiol Nutr Metab 51, 1–9.

Lulic-Kuryllo T & Inglis JG (2022). Sex differences in motor unit behaviour: A review. J Electromyogr Kinesiol 66, 102689.

Maharaj JN, Cresswell AG & Lichtwark GA (2019). Tibialis anterior tendinous tissue plays a key role in energy absorption during human walking. J Exp Biol 222, 1–7.

Marateb HR, McGill KC, Holobar A, Lateva ZC, Mansourian M & Merletti R (2011). Accuracy assessment of CKC high-density surface EMG decomposition in biceps femoris muscle. J Neural Eng 8, 066002.

Martinez-Valdes E, Negro F, Falla D, De Nunzio AM & Farina D (2018). Surface electromyographic amplitude does not identify differences in neural drive to synergistic muscles. J Appl Physiol 124, 1071–1079.

McGill KC & Dorfman LJ (1985). Automatic decomposition electromyography (ADEMG): validation and normative data in brachial biceps. Electroencephalogr Clin Neurophysiol 61, 453–461.

Merletti R & Farina D eds. (2016). Surface Electromyography : Physiology, Engineering, and Applications, 1st edn. Wiley. Available at: https://onlinelibrary.wiley.com/doi/book/10.1002/9781119082934 [Accessed February 11, 2026].

Navacchia A, Myers CA, Rullkoetter PJ & Shelburne KB (2016). Prediction of in vivo knee joint loads using a global probabilistic analysis. J Biomech Eng 138, 031002.

Negro F, Muceli S, Castronovo AM, Holobar A & Farina D (2016). Multi-channel intramuscular and surface EMG decomposition by convolutive blind source separation. J Neural Eng 13, 1–17.

Oskouei AE & Herzog W (2005). Observations on force enhancement in submaximal voluntary contractions of human adductor pollicis muscle. J Appl Physiol 98, 2087–2095.

Oskouei AE & Herzog W (2006). Force enhancement at different levels of voluntary contraction in human adductor pollicis. Eur J Appl Physiol 97, 280–287.

Pandy MG, Zajac FE, Sim E & Levine WS (1990). An optimal control model for maximum-height human jumping. J Biomech 23, 1185–1198.

Paquin J & Power GA (2018). History dependence of the EMG-torque relationship. J Electromyogr Kinesiol 41, 109–115.

Pinniger GJ & Cresswell AG (2007). Residual force enhancement after lengthening is present during submaximal plantar flexion and dorsiflexion actions in humans. J Appl Physiol 102, 18–25.

Pustejovsky JE, Pekofsky S & Zhang J (2025). clubSandwich: Cluster-Robust (Sandwich) Variance Estimators with Small-Sample Corrections. Available at: https://cran.r-project.org/web/packages/clubSandwich/index.html [Accessed January 6, 2026].

Raiteri BJ, Lauret L & Hahn D (2023). The force-length relation of the young adult human tibialis anterior. PeerJ 11, 1–23.

Raiteri BJ, Lauret L & Hahn D (2024). Residual force depression is not related to positive muscle fascicle work during submaximal voluntary dorsiflexion contractions in humans. J Physiol 602, 1085–1103.

Rousanoglou EN, Oskouei AE & Herzog W (2007). Force depression following muscle shortening in sub-maximal voluntary contractions of human adductor pollicis. J Biomech 40, 1–8.

Seiberl W, Hahn D, Herzog W & Schwirtz A (2012). Feedback controlled force enhancement and activation reduction of voluntarily activated quadriceps femoris during sub-maximal muscle action. J Electromyogr Kinesiol 22, 117–123.

Stoffel MA, Nakagawa S & Schielzeth H (2021). partR2 : partitioning R^2^ in generalized linear mixed models. PeerJ 9, e11414.

Taylor CA, Kopicko BH, Negro F & Thompson CK (2022). Sex differences in the detection of motor unit action potentials identified using high-density surface electromyography. J Electromyogr Kinesiol 65, 102675.

Tecchio P, Raiteri BJ & Hahn D (2024). Eccentric exercise ≠ eccentric contraction. J Appl Physiol 136, 954–965.

Trecarten N, Minozzo FC, Leite FS & Rassier DE (2015). Residual force depression in single sarcomeres is abolished by MgADP-induced activation. Sci Rep 5, 10555.

Van Noten P & Van Leemputte M (2011). The effect of muscle length on force depression after active shortening in soleus muscle of mice. Eur J Appl Physiol 111, 1361–1367.

Winter DA (2009). Biomechanics and motor control of human movement. John Wiley & Sons, Inc., Hoboken, NJ, USA. Available at: http://doi.wiley.com/10.1002/9780470549148 [Accessed August 31, 2022].

Yacyshyn AF, Mohammadalinejad G, Afsharipour B, Duchcherer J, Bashuk J, Bennett DJ, Negro F, Quinlan KA & Gorassini MA (2025). Sex-related differences in motoneuron firing behavior during typical development. J Neurophysiol 133, 1307–1319.

Zajac FE (1989). Muscle and tendon: properties, models, scaling, and application to biomechanics and motor control. Crit Rev Biomed Eng 17, 359–411.

