## Supporting information for "Stretch versus shortening contractions subsequently decrease versus increase neural drive to the human tibialis anterior"

### Agreement between decomposition approaches

*Rate of agreement*

For each motor unit shared between DEMUSE and MUedit at one contraction intensity, the rate of agreement (RoA) between approaches (i.e. manual editing for DEMUSE decompositions and automatic editing for MUedit decompositions) was calculated as described in Hug et al. (2021). A maximal lag of 50 ms was allowed between motor unit discharge patterns to find the best overlap, and a ±1 time point discharge time tolerance was used (i.e. discharge times were considered to agree if they did not differ by more than 0.5 ms).

A total of 456 RoA values were calculated (i.e. 2 contraction intensities × 4-30 common motor units). RoA decreased with contraction intensity, with mean (standard deviation) values of 82.0 (2.5)% and 80.3 (1.6)% at 20 and 40% MVT, respectively. RoA ranged from 43.7 to 97.2%, with 294 out of 456 values (64.5%) being lower than 85%. These RoA values are low compared with previous research (Hug *et al.*, 2021), indicating relatively poor agreement in discharge rate timings between the two approaches used in this study. Notably though, the RoA relative to the automatic editing approach increased following manual editing in DEMUSE compared with unedited DEMUSE decompositions (min-max: 28.9 to 96.3%; 75.8 (3.9)% and 74.8 (3.6)% at 20 and 40% MVT, respectively; Fig. S1). In line with previous research (Murks *et al.*, 2025), the number of common motor units also decreased with increasing contraction level (15 (6) and 12 (5) at 20 and 40% MVT, respectively) and 13 (5) common motor units were found across contraction intensities.

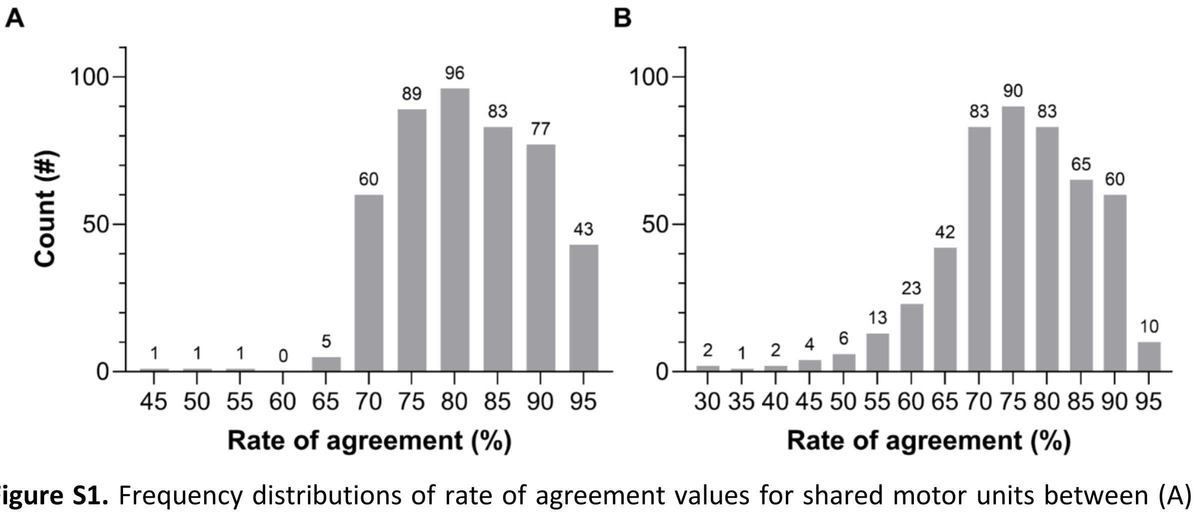


**Figure S1.** Frequency distributions of rate of agreement values for shared motor units between (A) manually edited DEMUSE decompositions and automatically edited MUedit decompositions and (B) unedited DEMUSE decompositions and automatically edited MUedit decompositions. Note that the automatic editing approach for MUedit decompositions led to a perfect rate of agreement relative to unedited MUedit decompositions. The mean rates of agreement were calculated by fitting linear mixed-effects models with a fixed contraction intensity effect (random intercepts and fixed slopes) without random motor unit effects nested under random participant effects (see the "Statistics" section within the manuscript for further details).

*Limits of agreement*

Limits of agreement between the mean discharge rates in each condition and at each contraction intensity between decomposition approaches was assessed via repeated-measures Bland-Altman analysis (note that the code was modified to fit linear mixed-effects models and use Satterthwaite's approximation to compute the denominator degrees of freedom for the t-statistics of the fixed-effects coefficients; Wade *et al.*, 2023).

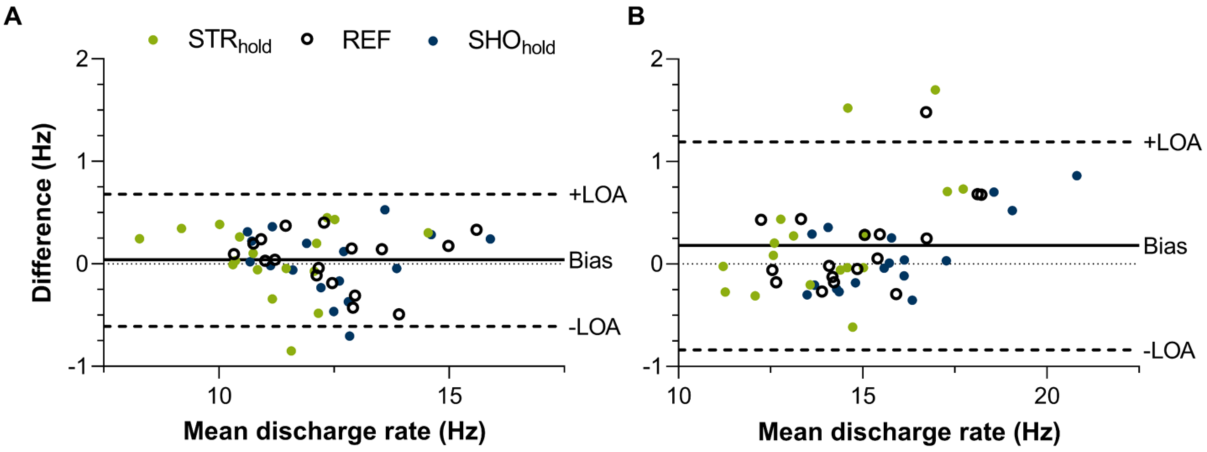


**Figure S2.** Agreement in mean discharge rates between manual editing of DEMUSE decompositions and automatic editing of MUedit decompositions at (A) 20% of maximum voluntary torque (MVT) and (B) 40% MVT based on repeated-measures Bland-Altman analysis. The agreement was high, with negligible bias (0 to 0.2 Hz) and relatively narrow 95% limits of agreement (LOA; maximum at 20% MVT: 0.7 Hz; maximum at 40% MVT: 1.2 Hz).

There was negligible systematic bias at 20 (0.04 (0.31) Hz) and 40% MVT (0.18 (0.48) Hz) between decomposition approaches, with random error being more prominent than systematic bias. The 95% limits of agreement were also narrow for both contraction intensities, but narrower at 20 (-0.61 to 0.68 Hz) than 40% MVT (-0.84 to 1.19 Hz; Fig. S2). These findings indicate high agreement in mean discharge rates between the two decomposition approaches used in this study. Consequently, due to the relatively minor differences between the manual and automatic editing approaches, the automatic editing approach may be favoured for calculating mean discharge rate during steady-state torque production (following visual verification of the motor unit pulse trains) due to its reduced subjectivity and time cost. Importantly though, the use of an automatic editing approach for assessing metrics requiring accurate discharge rate timings and for calculating mean discharge rates when force varies remains questionable. In such cases, manual editing based on strict, well-defined rules incorporating muscle physiology would be preferable (e.g., Del Vecchio *et al.*, 2020; Hug *et al.*, 2021).

### Decomposition settings

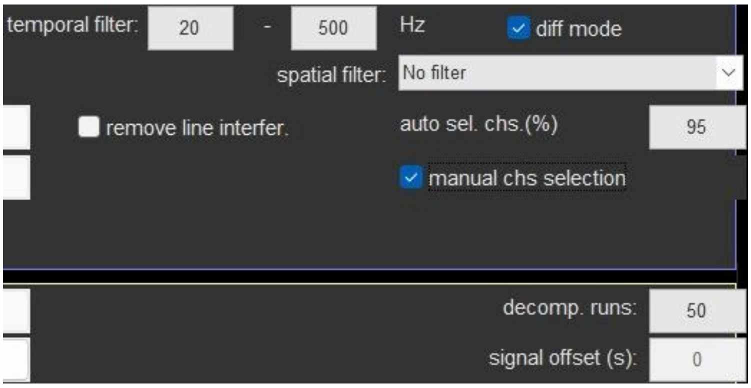


**Figure S3.** Screenshot of the DEMUSE (v5.01; Holobar & Zazula, 2007*a*, 2007*b*) decomposition settings used in MATLAB, with a default decomposition speed of "slower and accurate". Other decomposition settings included: Smart PT orthogonalization = on; Sparse decomposition = off; Minimal number of motor unit firings = 10; Maximal coefficient of variation of motor unit inter-spike interval = 1.5; Pulse height = 1.2. DEMUSE filtered the high-density surface electromyography signals with a bandpass filter between 20 and 500 Hz using a dual-pass fourth-order Butterworth filter. The "diff mode" option was selected to enhance local signal components and suppress common-mode noise by converting monopolar signals into adjacent-row (longitudinal) single-differential signals. 5% of the high-density surface electromyography channels (i.e. 3 channels) were then removed based on the lowest ranking correlation coefficients between each channel and its immediate neighbours, which is a standard preprocessing step in DEMUSE before decomposition. Following decomposition, the motor unit discharge patterns were manually edited according to previous guidelines (Del Vecchio *et al.*, 2020; Hug *et al.*, 2021) and duplicate motor units were removed if two motor units shared more than 30% of their identified discharge times (within a time interval of 0.5 ms), with the lower pulse-to-noise ratio motor unit being excluded (Holobar *et al.*, 2010). Subsequently, the median motor unit discharge rate was calculated for the matched manually edited motor units among conditions over condition-specific intervals to be consistent with the median calculation used as part of the automatic approach with MUedit (see Fig. S4 caption below).

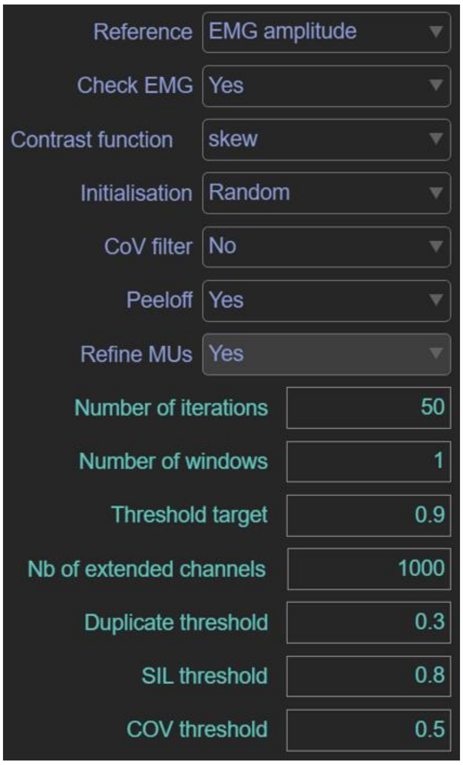


**Figure S4.** Screenshot of the MUedit (v1.2; Avrillon *et al.*, 2024) decomposition settings used in MATLAB. By default, MUedit filters the high-density surface electromyography signals with an adaptive notch filter to remove frequencies with abnormal peaks, as well as a bandpass filter between 20 and 500 Hz using a dual-pass second-order Butterworth filter. A "random" initialisation was used based on the description in Avrillon et al. (2024) that random weights with the function "skew" resulted in slightly better performance than with an initialization with the maximal EMG values ("EMG max"). Following decomposition with MUedit, duplicate motor units were removed if there were more than 30% of shared discharge times (within a time interval of 0.5 ms) between two motor units, with the lower pulse-to-noise ratio motor unit being excluded (Holobar *et al.*, 2010). Subsequently, the motor unit discharge patterns were automatically edited (see the "Decomposition editing" section within the manuscript) and visually verified before the median motor unit discharge rate was calculated for the matched automatically edited motor units among conditions over condition-specific intervals to avoid outliers from affecting the central tendency.

### Active torque model fitting and results

Fixed effects were fitted using a linear mixed-effects model. Model-based standard errors and p-values are presented. Sandwich variance estimation-based standard errors and p-values were also produced; these do not assume normality of the data. A Wald statistic based on the sandwich variance estimate was also produced to test the effect of factor condition.

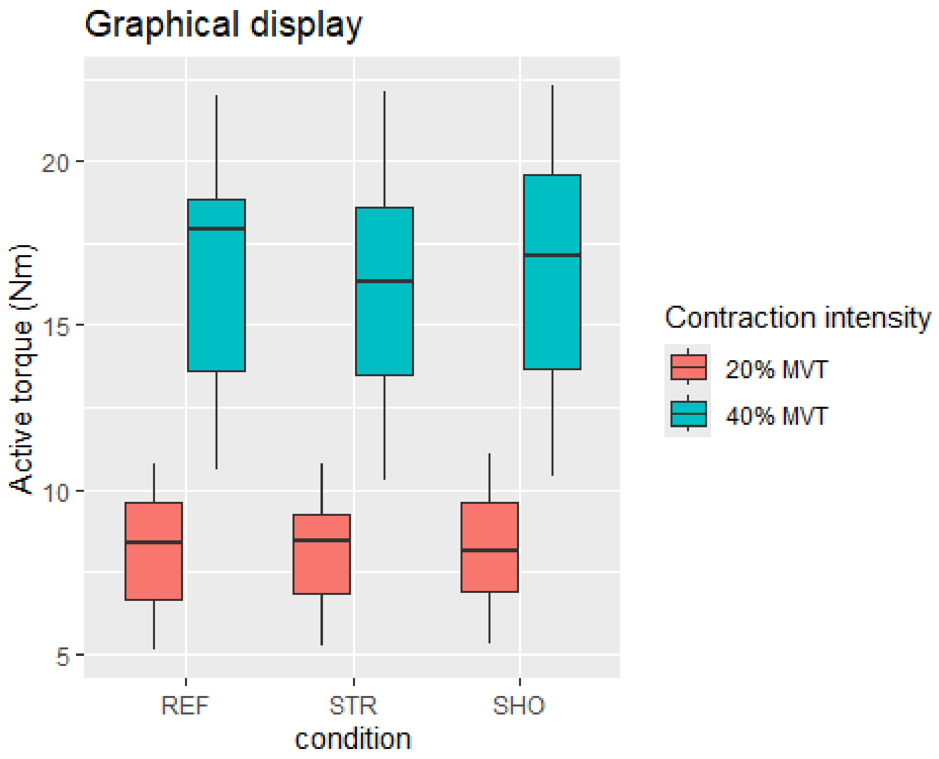


*Fixed-effects model with no interaction and with/without random trial effects nested under random participant effects*

This fixed-effects model fits the factors intensity and condition with random intercepts and fixed slopes. We first compared fitting the model by REML vs. ML, and ML resulted in a better fit without random trial effects nested under random participant effects:

```
## $statistics
##                         aic     bic bayes.factor
## no_interaction_REML 894.617 916.698        0.046
## no_interaction_ML   888.461 910.542       21.718
##
## $predicted_differences
##    0%   25%   50%   75%  100%
## 0.000 0.001 0.001 0.001 0.001

## $statistics
##                             aic     bic bayes.factor
## no_interaction_ML       888.461 910.542       17.117
## no_interaction_ML_trial 890.461 916.222        0.058
```

```
##
## $predicted_differences
##   0%  25%  50%  75% 100%
##    0    0    0    0    0
```

*Model summary*

```
## Linear mixed model fit by maximum likelihood . t-tests use Satterthwait
e's
##   method [lmerModLmerTest]
## Formula: outcome ~ intensity + condition + (1 | pid)
##    Data: d
##
##      AIC      BIC   logLik deviance df.resid
##    888.5    910.5   -438.2    876.5      287
##
## Scaled residuals:
##      Min       1Q   Median       3Q      Max
## -2.03677 -0.70783  0.06513  0.71736  1.75571
##
## Random effects:
##  Groups   Name        Variance Std.Dev.
##  pid      (Intercept) 7.405    2.7212
##  Residual             0.873    0.9343
## Number of obs: 293, groups:  pid, 17
##
## Fixed effects:
##             Estimate Std. Error       df t value Pr(>|t|)
## (Intercept)   8.1670     0.6692  17.7365  12.203 4.62e-10 ***
## intensity2    8.0349     0.1098 276.0476  73.189  < 2e-16 ***
## condition2   -0.1531     0.1351 276.0497  -1.133    0.258
## condition3    0.0217     0.1362 276.0517   0.159    0.874
## ---
## Signif. codes:  0 '***' 0.001 '**' 0.01 '*' 0.05 '.' 0.1 ' ' 1
##
## Correlation of Fixed Effects:
##            (Intr) intns2 cndtn2
## intensity2 -0.078
## condition2 -0.105 -0.018
## condition3 -0.102 -0.038  0.523
```

*Model- & sandwich-based standard errors and p-values*

The full cluster-robust variance-covariance matrix was calculated, which includes the variances (SEs) and covariances (how much the estimates correlate with each other):

```
## Model- & sandwich-based standard errors and p-values

## Click on the arrow for sandwich-based p-values.

## # A tibble: 4 × 6
##   term        estimate `Model-based s.e.` `Model-based p-value` `Sandwi
ch s.e.`
##   <chr>          <dbl>              <dbl>                 <dbl>
<dbl>
## 1 (Intercept)   8.17               0.669             7.14e- 28
0.442
## 2 intensity2    8.03               0.110             1.10e-187
0.464
## 3 condition2   -0.153              0.135             2.58e-  1
0.0670
## 4 condition3    0.0217             0.136             8.74e-  1
0.0748
##   `Sandwich p-value`
##              <dbl>
## 1           3.27e-12
## 2           1.08e-11
## 3           3.65e- 2
## 4           7.76e- 1
```

*Fixed-effects model with interaction*

This fixed-effects model fits intensity and condition and the interaction term intensity:condition with random intercepts and fixed slopes. The fixed-effects model with an intensity:condition interaction did not result in a better fit:

```
## $statistics
##                        aic     bic bayes.factor
## no_interaction_ML 888.461 910.542      278.323
## interaction_ML    892.358 921.799        0.004
##
## $predicted_differences
##    0%   25%   50%   75%  100%
## 0.003 0.008 0.016 0.024 0.031
```

*Random-effects model with no interaction*

This random-effects model fits intensity and condition with random intercepts and random slopes. The random-effects model with no interaction resulted in a better fit:

```
## $statistics
##                                aic     bic  bayes.factor
## no_interaction_ML         888.461 910.542  0.000000e+00
## no_interaction_ML_random -139.523 -84.320 1.075357e+216
##
## $predicted_differences
##    0%   25%   50%   75%  100%
## 0.028 0.503 0.628 1.201 1.783
```

*Model summary*

```
## Linear mixed model fit by maximum likelihood . t-tests use Satterthwait
e's
##   method [lmerModLmerTest]
## Formula: outcome ~ intensity + condition + (intensity + condition | pid
)
##    Data: d
##
##      AIC      BIC   logLik deviance df.resid
##   -139.5    -84.3     84.8   -169.5      278
##
## Scaled residuals:
##     Min      1Q  Median      3Q     Max
## -4.0205 -0.4728  0.0082  0.4435  3.7984
##
## Random effects:
##  Groups   Name        Variance Std.Dev. Corr
##  pid      (Intercept) 3.25858  1.8052
##           intensity2  3.10115  1.7610    0.99
##           condition2  0.02476  0.1574   -0.05 -0.05
##           condition3  0.01149  0.1072    0.16  0.22  0.83
##  Residual             0.01550  0.1245
## Number of obs: 293, groups:  pid, 17
##
## Fixed effects:
##             Estimate Std. Error       df t value Pr(>|t|)
## (Intercept)  8.11256    0.43807 17.00994  18.519 1.04e-12 ***
## intensity2   8.04477    0.42736 17.00936  18.824 7.95e-13 ***
## condition2  -0.09038    0.04226 17.05032  -2.139   0.0472 *
## condition3   0.04707    0.03179 17.28184   1.481   0.1566
## ---
## Signif. codes:  0 '***' 0.001 '**' 0.01 '*' 0.05 '.' 0.1 ' ' 1
##
## Correlation of Fixed Effects:
##            (Intr) intns2 cndtn2
## intensity2  0.990
## condition2 -0.053 -0.047
## condition3  0.121  0.175  0.744
```

*Model- & sandwich-based standard errors and p-values*

```
## Model- & sandwich-based standard errors and p-values

## Click on the arrow for sandwich-based p-values.

## # A tibble: 4 × 6
##   term        estimate `Model-based s.e.` `Model-based p-value` `Sandwi
ch s.e.`
##   <chr>          <dbl>              <dbl>                 <dbl>
<dbl>
## 1 (Intercept)   8.11               0.438                 1.97e-50
0.452
## 2 intensity2    8.04               0.427                 1.56e-51
0.441
## 3 condition2   -0.0904             0.0423                3.33e- 2
0.0435
## 4 condition3    0.0471             0.0318                1.40e- 1
0.0328
##   `Sandwich p-value`
##                <dbl>
## 1           4.98e-12
## 2           3.88e-12
## 3           5.44e- 2
## 4           1.70e- 1
```

*Estimated marginal means*

Estimated marginal means (EMMs) for the random-effects model with no interaction are reported below.

```
##  intensity emmean    SE df lower.CL upper.CL
##  1            8.1 0.453 17     7.14     9.05
##  2           16.1 0.891 17    14.26    18.02
##
## Results are averaged over the levels of: condition
## Degrees-of-freedom method: satterthwaite
## Confidence level used: 0.95

##  condition emmean    SE df lower.CL upper.CL
##  1           12.1 0.670 17     10.7     13.5
##  2           12.0 0.670 17     10.6     13.5
##  3           12.2 0.676 17     10.8     13.6
##
## Results are averaged over the levels of: intensity
## Degrees-of-freedom method: satterthwaite
## Confidence level used: 0.95
```

### Coefficient of variation in active torque (torque steadiness) model fitting and results

Fixed effects were fitted using a linear mixed-effects model. Model-based standard errors and p-values are presented. Sandwich variance estimation-based standard errors and p-values were also produced; these do not assume normality of the data. A Wald statistic based on the sandwich variance estimate was also produced to test the effect of factor condition.

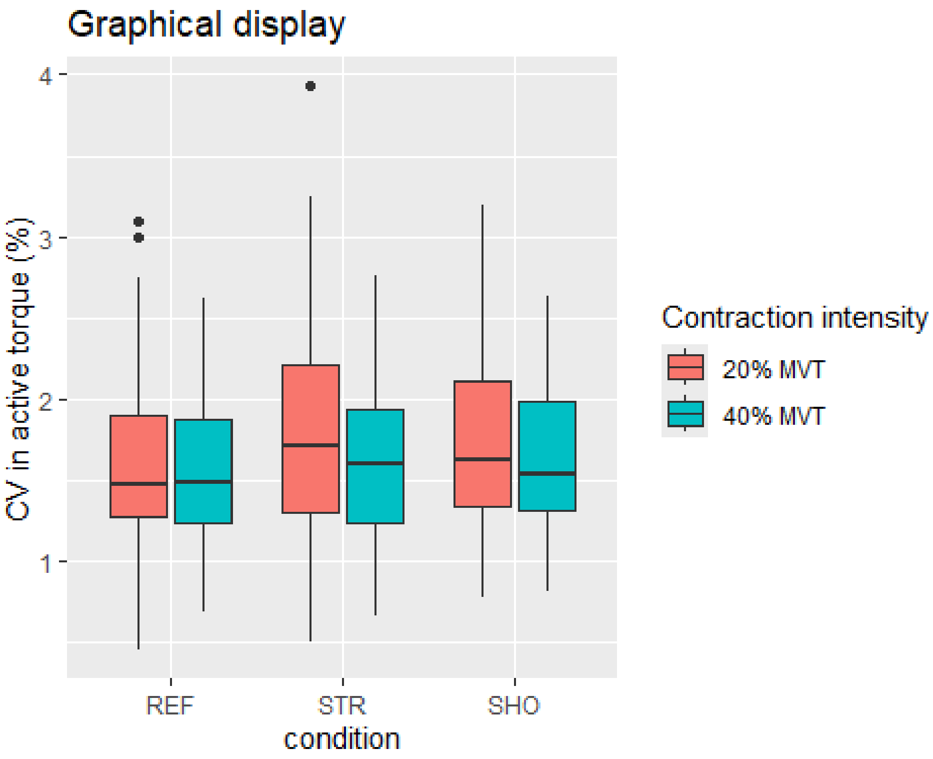


*Fixed-effects model with no interaction and with/without random trial effects nested under random participant effects*

This fixed-effects model fits the factors intensity and condition with random intercepts and fixed slopes. We first compared fitting the model by REML vs. ML, and ML resulted in a better fit´without random trial effects nested under random participant effects:

```
## $statistics
##                         aic     bic bayes.factor
## no_interaction_REML 460.823 482.904        0.001
## no_interaction_ML   446.371 468.452     1374.160
##
## $predicted_differences
##    0%   25%   50%   75%  100%
## 0.000 0.000 0.001 0.002 0.005

## $statistics
##                               aic     bic bayes.factor
## no_interaction_ML       446.371 468.452       17.117
## no_interaction_ML_trial 448.371 474.133        0.058
```

```
##
## $predicted_differences
##   0%  25%  50%  75% 100%
##    0    0    0    0    0
```

*Model summary*

```
## Linear mixed model fit by maximum likelihood . t-tests use Satterthwait
e's
##   method [lmerModLmerTest]
## Formula: outcome ~ intensity + condition + (1 | pid)
##    Data: d
##
##      AIC      BIC   logLik deviance df.resid
##    446.4    468.5   -217.2    434.4      287
##
## Scaled residuals:
##     Min      1Q  Median      3Q     Max
## -2.5177 -0.6375 -0.0702  0.5561  3.1991
##
## Random effects:
##  Groups   Name        Variance Std.Dev.
##  pid      (Intercept) 0.09018  0.3003
##  Residual             0.22896  0.4785
## Number of obs: 293, groups:  pid, 17
##
## Fixed effects:
##              Estimate Std. Error        df t value Pr(>|t|)
## (Intercept)   1.66414    0.09233  32.63114  18.023   <2e-16 ***
## intensity2   -0.12294    0.05619 276.54477  -2.188   0.0295 *
## condition2    0.08490    0.06915 276.59870   1.228   0.2206
## condition3    0.09131    0.06970 276.63483   1.310   0.1912
## ---
## Signif. codes:  0 '***' 0.001 '**' 0.01 '*' 0.05 '.' 0.1 ' ' 1
##
## Correlation of Fixed Effects:
##            (Intr) intns2 cndtn2
## intensity2 -0.290
## condition2 -0.388 -0.018
## condition3 -0.378 -0.038  0.523
```

*Model- & sandwich-based standard errors and p-values*

The full cluster-robust variance-covariance matrix was calculated, which includes the variances (SEs) and covariances (how much the estimates correlate with each other):

```
## Model- & sandwich-based standard errors and p-values

## Click on the arrow for sandwich-based p-values.

## # A tibble: 4 × 6
##   term        estimate `Model-based s.e.` `Model-based p-value` `Sandwi
ch s.e.`
##   <chr>          <dbl>              <dbl>                 <dbl>
<dbl>
## 1 (Intercept)   1.66               0.0923              4.30e-49
0.130
## 2 intensity2   -0.123              0.0562              2.95e- 2
0.148
## 3 condition2    0.0849             0.0691              2.21e- 1
0.0496
## 4 condition3    0.0913             0.0697              1.91e- 1
0.0549
##   `Sandwich p-value`
##              <dbl>
## 1          8.00e-10
## 2          4.18e- 1
## 3          1.07e- 1
## 4          1.16e- 1
```

*Fixed-effects model with interaction*

This fixed-effects model fits intensity and condition and the interaction term intensity:condition with random intercepts and fixed slopes. The fixed-effects model with an intensity:condition interaction did not result in a better fit:

```
## $statistics
##                       aic     bic bayes.factor
## no_interaction_ML 446.371 468.452      133.201
## interaction_ML    448.795 478.236        0.008
##
## $predicted_differences
##    0%   25%   50%   75%  100%
## 0.014 0.022 0.032 0.045 0.052
```

*Random-effects model with no interaction*

This random-effects model fits intensity and condition with random intercepts and random slopes. The random-effects model with no interaction resulted in a better fit:

```
## $statistics
##                              aic     bic bayes.factor
## no_interaction_ML        446.371 468.452            0
## no_interaction_ML_random 380.198 435.400     15037364
##
## $predicted_differences
##    0%   25%   50%   75%  100%
## 0.003 0.095 0.199 0.308 0.621
```

*Model summary*

```
## Linear mixed model fit by maximum likelihood . t-tests use Satterthwait
e's
##   method [lmerModLmerTest]
## Formula: outcome ~ intensity + condition + (intensity + condition | pid
)
##    Data: d
##
##      AIC      BIC   logLik deviance df.resid
##    380.2    435.4   -175.1    350.2      278
##
## Scaled residuals:
##     Min      1Q  Median      3Q     Max
## -2.5945 -0.6436 -0.1209  0.6019  3.9019
##
## Random effects:
##  Groups   Name        Variance  Std.Dev. Corr
##  pid      (Intercept) 0.2269692 0.47641
##           intensity2  0.3019572 0.54951  -0.83
##           condition2  0.0007678 0.02771   0.11  0.47
##           condition3  0.0018114 0.04256   0.92 -0.98 -0.29
##  Residual             0.1482100 0.38498
## Number of obs: 293, groups:  pid, 17
##
## Fixed effects:
##              Estimate Std. Error        df t value Pr(>|t|)
## (Intercept)   1.67477    0.12429  17.23823  13.475 1.39e-10 ***
## intensity2   -0.14225    0.14081  17.00624  -1.010    0.327
## condition2    0.08328    0.05630 178.38248   1.479    0.141
## condition3    0.08002    0.05729 129.05304   1.397    0.165
## ---
## Signif. codes:  0 '***' 0.001 '**' 0.01 '*' 0.05 '.' 0.1 ' ' 1
##
## Correlation of Fixed Effects:
##            (Intr) intns2 cndtn2
## intensity2 -0.784
## condition2 -0.219  0.047
## condition3 -0.068 -0.180  0.506
## optimizer (nloptwrap) convergence code: 0 (OK)
## boundary (singular) fit: see help('isSingular')
```

*Model- & sandwich-based standard errors and p-values*

```
## Model- & sandwich-based standard errors and p-values

## Click on the arrow for sandwich-based p-values.

## # A tibble: 4 × 6
##   term        estimate `Model-based s.e.` `Model-based p-value` `Sandwi
ch s.e.`
##   <chr>          <dbl>              <dbl>                 <dbl>
<dbl>
## 1 (Intercept)   1.67               0.124               3.42e-32
0.127
## 2 intensity2   -0.142              0.141               3.13e- 1
0.145
## 3 condition2    0.0833             0.0563              1.40e- 1
0.0461
## 4 condition3    0.0800             0.0573              1.64e- 1
0.0564
##   `Sandwich p-value`
##                <dbl>
## 1           5.29e-10
## 2           3.42e- 1
## 3           8.99e- 2
## 4           1.75e- 1
```

*Estimated marginal means*

Estimated marginal means (EMMs) for the random-effects model with no interaction are reported below.

```
##  intensity emmean     SE   df lower.CL upper.CL
##  1           1.73 0.1270 17.0     1.46     2.00
##  2           1.59 0.0852 16.6     1.41     1.77
##
## Results are averaged over the levels of: condition
## Degrees-of-freedom method: satterthwaite
## Confidence level used: 0.95

##  condition emmean     SE   df lower.CL upper.CL
##  1           1.60 0.0825 17.8     1.43     1.78
##  2           1.69 0.0850 17.2     1.51     1.87
##  3           1.68 0.0889 16.8     1.50     1.87
##
## Results are averaged over the levels of: intensity
## Degrees-of-freedom method: satterthwaite
## Confidence level used: 0.95
```

### Normalised global EMG amplitude model fitting and results

Fixed effects were fitted using a linear mixed-effects model. Model-based standard errors and p-values are presented. Sandwich variance estimation-based standard errors and p-values were also produced; these do not assume normality of the data. A Wald statistic based on the sandwich variance estimate was also produced to test the effect of factor condition.

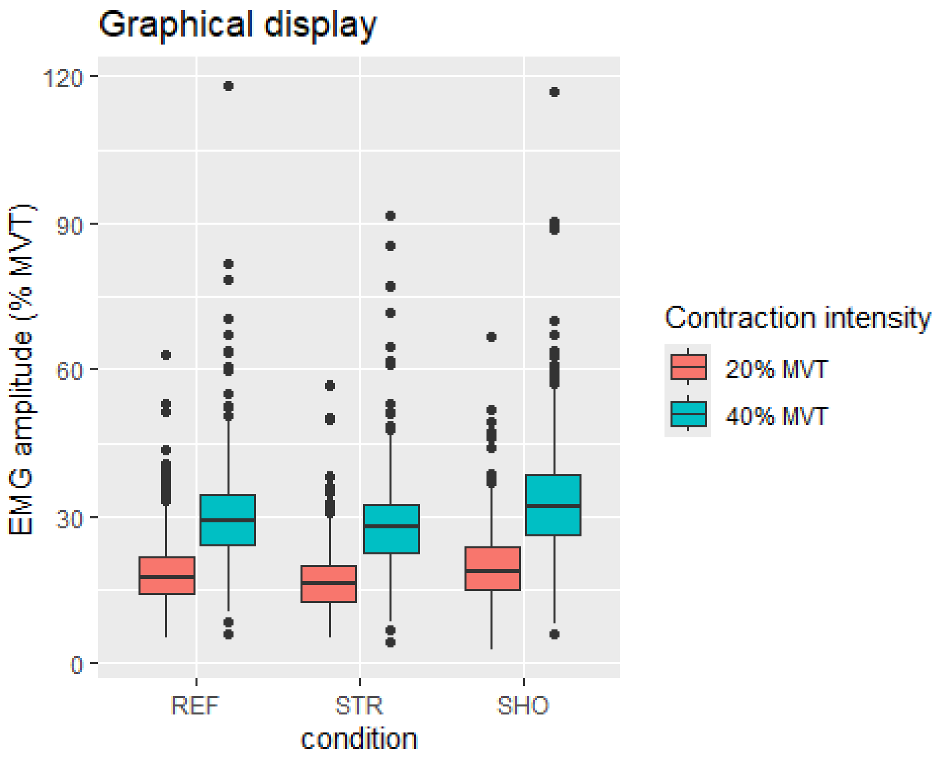


*Fixed-effects model with no interaction and with/without random trial effects nested under random participant effects*

This fixed-effects model fits the factors intensity and condition with random intercepts and fixed slopes. We first compared fitting the model by REML vs. ML, and the models were similar, but not equivalent, with ML resulting in the better fit. The model with random channel effects nested under random participant effects resulted in a better fit vs. a model with random participant effects only:

```
## $statistics
##                          aic      bic  bayes.factor
## no_interaction_REML 38865.71 38905.92         0.236
## no_interaction_ML   38862.82 38903.03         4.245
##
## $predicted_differences
##    0%    25%    50%    75%   100%
## 0.000 0.001 0.001 0.001 0.002
```

```
## $statistics
##                                 aic      bic bayes.factor
## no_interaction_ML         38862.82 38903.03            0
## no_interaction_ML_channel 33875.84 33922.76          Inf
##
## $predicted_differences
##     0%    25%    50%    75%   100%
##  0.001  1.213  2.625  4.363 54.565
```

*Model summary*

```
## Linear mixed model fit by maximum likelihood . t-tests use Satterthwait
e's
##   method [lmerModLmerTest]
## Formula: outcome ~ intensity + condition + (1 | pid/channel)
##    Data: d
##
##      AIC      BIC   logLik deviance df.resid
##  33875.8  33922.8 -16930.9  33861.8     6011
##
## Scaled residuals:
##     Min      1Q  Median      3Q     Max
## -9.3069 -0.5464 -0.0679  0.4849  9.4813
##
## Random effects:
##  Groups      Name        Variance Std.Dev.
##  channel:pid (Intercept) 27.06    5.202
##  pid         (Intercept) 23.50    4.848
##  Residual                10.01    3.164
## Number of obs: 6018, groups:  channel:pid, 1003; pid, 17
##
## Fixed effects:
##              Estimate Std. Error         df t value Pr(>|t|)
## (Intercept)   18.07228    1.18992   17.12037   15.19 2.28e-11 ***
## intensity2    12.19226    0.08156 5015.00312  149.48  < 2e-16 ***
## condition2    -1.63893    0.09990 5015.00315  -16.41  < 2e-16 ***
## condition3     2.22589    0.09990 5015.00315   22.28  < 2e-16 ***
## ---
## Signif. codes:  0 '***' 0.001 '**' 0.01 '*' 0.05 '.' 0.1 ' ' 1
##
## Correlation of Fixed Effects:
##            (Intr) intns2 cndtn2
## intensity2 -0.034
## condition2 -0.042  0.000
## condition3 -0.042  0.000  0.500
```

*Model- & sandwich-based standard errors and p-values*

The full cluster-robust variance-covariance matrix was calculated, which includes the variances (SEs) and covariances (how much the estimates correlate with each other):

```
## Model- & sandwich-based standard errors and p-values

## Click on the arrow for sandwich-based p-values.

## # A tibble: 4 × 6
##   term        estimate `Model-based s.e.` `Model-based p-value` `Sandwi
ch s.e.`
##   <chr>          <dbl>              <dbl>                 <dbl>
<dbl>
## 1 (Intercept)    18.1              1.19                 3.75e- 51
1.10
## 2 intensity2     12.2              0.0816               0
0.623
## 3 condition2     -1.64             0.0999               3.28e- 59
0.422
## 4 condition3      2.23             0.0999               9.54e-106
0.436
##   `Sandwich p-value`
##                <dbl>
## 1           1.89e-11
## 2           1.32e-12
## 3           1.31e- 3
## 4           1.06e- 4
```

*Condition p-value*

The Wald statistic testing for an effect of condition is:

```
##  test Fstat df_num df_denom     p_val sig
##   HTZ  18.1      2       15 0.0001012 ***
```

*Post-hoc comparisons*

Estimated marginal means (EMMs) for the fixed-effects model with no interaction, while explicitly using cluster-robust standard errors (which accounts for non-independent data or heteroskedasticity), are reported below. This approach was used to ensure that the confidence intervals and subsequent p-values for the EMMs are more accurate and reliable.

```
##  intensity condition emmean   SE   df lower.CL upper.CL
##  1         1           18.1 1.10 17.1     15.8     20.4
##  2         1           30.3 1.43 17.1     27.2     33.3
##  1         2           16.4 1.03 17.1     14.3     18.6
##  2         2           28.6 1.30 17.1     25.9     31.4
##  1         3           20.3 1.21 17.1     17.7     22.9
##  2         3           32.5 1.62 17.1     29.1     35.9
##
## Degrees-of-freedom method: satterthwaite
## Confidence level used: 0.95

## condition = 1:
##  contrast                estimate    SE   df t.ratio p.value
##  intensity1 - intensity2    -12.2 0.623 5015 -19.585  <.0001
##
```

```
## condition = 2:
##  contrast                 estimate    SE   df t.ratio p.value
##  intensity1 - intensity2    -12.2 0.623 5015 -19.585  <.0001
##
## condition = 3:
##  contrast                 estimate    SE   df t.ratio p.value
##  intensity1 - intensity2    -12.2 0.623 5015 -19.585  <.0001
##
## Degrees-of-freedom method: satterthwaite
```

```
## intensity = 1:
##  contrast                 estimate    SE   df t.ratio p.value
##  condition1 - condition2     1.64 0.422 5015   3.886  0.0001
##  condition1 - condition3    -2.23 0.436 5015  -5.103  <.0001
##  condition2 - condition3    -3.86 0.628 5015  -6.157  <.0001
##
## intensity = 2:
##  contrast                 estimate    SE   df t.ratio p.value
##  condition1 - condition2     1.64 0.422 5015   3.886  0.0001
##  condition1 - condition3    -2.23 0.436 5015  -5.103  <.0001
##  condition2 - condition3    -3.86 0.628 5015  -6.157  <.0001
##
## Degrees-of-freedom method: satterthwaite
## P value adjustment: holm method for 3 tests
```

*Fixed-effects model with interaction*

This fixed-effects model fits intensity and condition and the interaction term intensity:condition with random intercepts and fixed slopes. The fixed-effects model with an intensity:condition interaction resulted in a better fit:

```
## $statistics
##                                 aic      bic bayes.factor
## no_interaction_ML_channel 33875.84 33922.76 0.000000e+00
## interaction_ML_channel    33725.99 33786.31 4.246506e+29
##
## $predicted_differences
##    0%   25%   50%   75%  100%
## 0.206 0.312 0.401 0.703 0.810
```

*Model summary*

```
## Linear mixed model fit by maximum likelihood . t-tests use Satterthwait
e's
##   method [lmerModLmerTest]
## Formula: outcome ~ intensity * condition + (1 | pid/channel)
##    Data: d
##
##      AIC      BIC   logLik deviance df.resid
##  33726.0  33786.3 -16854.0  33708.0     6009
##
## Scaled residuals:
##     Min      1Q  Median      3Q     Max
## -9.2449 -0.5198 -0.0324  0.4769  9.6938
```

```
##
## Random effects:
##  Groups      Name        Variance Std.Dev.
##  channel:pid (Intercept) 27.115   5.207
##  pid         (Intercept) 23.498   4.848
##  Residual                 9.707   3.116
## Number of obs: 6018, groups:  channel:pid, 1003; pid, 17
##
## Fixed effects:
##                          Estimate Std. Error        df t value Pr(>|t|)
## (Intercept)               18.3795     1.1912   17.1949  15.429 1.66e-11 *
**
## intensity2                11.5778     0.1391 5015.0002  83.220  < 2e-16 *
**
## condition2                -1.5455     0.1391 5015.0002 -11.109  < 2e-16 *
**
## condition3                 1.2108     0.1391 5015.0002   8.703  < 2e-16 *
**
## intensity2:condition2     -0.1869     0.1967 5015.0002  -0.950    0.342
## intensity2:condition3      2.0302     0.1967 5015.0002  10.318  < 2e-16 *
**
## ---
## Signif. codes:  0 '***' 0.001 '**' 0.01 '*' 0.05 '.' 0.1 ' ' 1
##
## Correlation of Fixed Effects:
##             (Intr) intns2 cndtn2 cndtn3 int2:2
## intensity2  -0.058
## condition2  -0.058  0.500
## condition3  -0.058  0.500  0.500
## intnsty2:c2  0.041 -0.707 -0.707 -0.354
## intnsty2:c3  0.041 -0.707 -0.354 -0.707  0.500
```

*Model- & sandwich-based standard errors and p-values*

```
## Model- & sandwich-based standard errors and p-values

## Click on the arrow for sandwich-based p-values.

## # A tibble: 6 × 6
##   term                  estimate `Model-based s.e.` `Model-based p-valu
e`
##   <chr>                    <dbl>              <dbl>                <db
l>
## 1 (Intercept)             18.4                1.19             1.05e-
52
## 2 intensity2              11.6                0.139            0
## 3 condition2              -1.55               0.139            2.15e-
28
## 4 condition3               1.21               0.139            4.12e-
18
## 5 intensity2:condition2   -0.187              0.197            3.42e-
1
## 6 intensity2:condition3    2.03               0.197            9.35e-
25
```

```
##   `Sandwich s.e.` `Sandwich p-value`
##             <dbl>              <dbl>
## 1           1.11            1.80e-11
## 2           0.753           5.31e-11
## 3           0.399           1.33e- 3
## 4           0.458           1.77e- 2
## 5           0.683           7.88e- 1
## 6           0.738           1.42e- 2
```

*Interaction, intensity & condition p-values*

The Wald statistics testing for an effect of the intensity:condition interaction term, for an effect of the intensity term and the intensity:condition interaction term, and for an effect of the condition term and the intensity:condition interaction term are:

```
##  test Fstat df_num df_denom   p_val sig
##   HTZ  6.22      2       15 0.01078   *

##  test Fstat df_num df_denom   p_val sig
##   HTZ   124      3       14 < 1e-04 ***

##  test Fstat df_num df_denom    p_val sig
##   HTZ  8.43      4       13 0.001396  **
```

*Post-hoc comparisons*

Estimated marginal means (EMMs) for the interaction between intensity and condition for the fixed-effects model, while explicitly using cluster-robust standard errors (which accounts for non-independent data or heteroskedasticity), are reported below. This approach was used to ensure that the confidence intervals and subsequent p-values for the EMMs are more accurate and reliable.

```
##  intensity condition emmean   SE   df lower.CL upper.CL
##  1         1           18.4 1.11 17.2     16.0     20.7
##  2         1           30.0 1.45 17.2     26.9     33.0
##  1         2           16.8 1.03 17.2     14.7     19.0
##  2         2           28.2 1.29 17.2     25.5     30.9
##  1         3           19.6 1.19 17.2     17.1     22.1
##  2         3           33.2 1.68 17.2     29.6     36.7
## 
## Degrees-of-freedom method: satterthwaite 
## Confidence level used: 0.95

## condition = 1:
##  contrast                  estimate    SE   df t.ratio p.value
##  intensity1 - intensity2    -11.6 0.753 5015 -15.369  <.0001
## 
## condition = 2:
##  contrast                  estimate    SE   df t.ratio p.value
##  intensity1 - intensity2    -11.4 0.585 5015 -19.475  <.0001
## 
```

```
## condition = 3:
##  contrast                 estimate    SE   df t.ratio p.value
##  intensity1 - intensity2    -13.6 0.852 5015 -15.966  <.0001
##
## Degrees-of-freedom method: satterthwaite
```

```
## intensity = 1:
##  contrast                 estimate    SE   df t.ratio p.value
##  condition1 - condition2     1.55 0.399 5015   3.878  0.0002
##  condition1 - condition3    -1.21 0.458 5015  -2.643  0.0082
##  condition2 - condition3    -2.76 0.593 5015  -4.648  <.0001
##
## intensity = 2:
##  contrast                 estimate    SE   df t.ratio p.value
##  condition1 - condition2     1.73 0.656 5015   2.642  0.0083
##  condition1 - condition3    -3.24 0.666 5015  -4.868  <.0001
##  condition2 - condition3    -4.97 0.801 5015  -6.213  <.0001
##
## Degrees-of-freedom method: satterthwaite
## P value adjustment: holm method for 3 tests
```

*Random-effects model with interaction*

This random-effects model fits intensity and condition and the interaction term intensity:condition with random intercepts and random slopes. The fixed-effects model with an intensity:condition interaction resulted in a better fit (note that a random-effects model with random channel effects nested under random participant effects could not be fit because the number of observations (6018) was less than or equal to the number of random effects for the term (intensity*condition|pid/channel):

```
## $statistics
##                              aic      bic bayes.factor
## interaction_ML_channel 33725.99 33786.31          Inf
## interaction_ML_random  38456.78 38644.45            0
##
## $predicted_differences
##     0%    25%    50%    75%   100%
##  0.001  1.305  2.790  4.703 56.670
```

### Motor unit discharge rate model fitting and results (assisted by Alain C. Vandal)

Fixed effects were fitted using a linear mixed-effects model with motor units as random effects nested under the participant id. Model-based standard errors and p-values are presented. Sandwich variance estimation-based standard errors and p-values were also produced; these do not assume normality of the data. A Wald statistic based on the sandwich variance estimate was also produced to test the effect of factor condition.

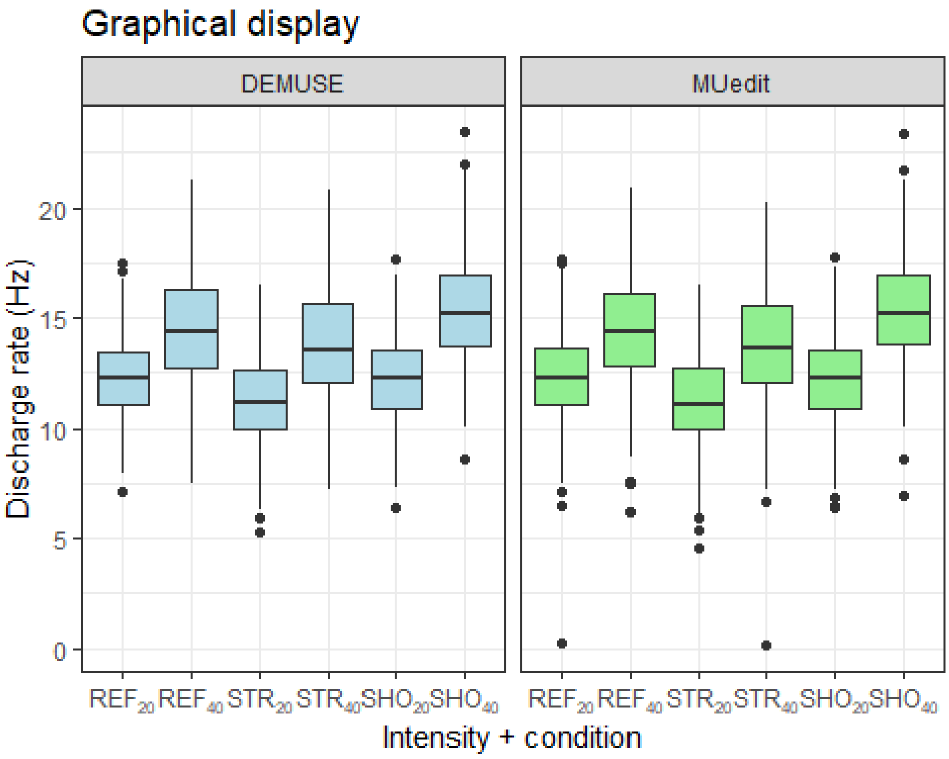


*Fixed-effects model with no interaction and with/without random motor unit effects nested under random participant effects*

This fixed-effects model fits the factors decomp, intensity, and condition with random intercepts and fixed slopes. We first compared fitting the model by REML vs. ML, and ML resulted in a better fit with random motor unit effects nested under random participant effects:

```
## $statistics
##                          aic      bic  bayes.factor
## no_interaction_REML 14967.73 15017.75         0.000
## no_interaction_ML   14951.80 15001.82      2878.726
##
## $predicted_differences
##    0%   25%   50%   75%  100%
## 0.000 0.000 0.000 0.001 0.004
```

```
## $statistics
##                              aic      bic bayes.factor
## no_interaction_ML     14951.80 15001.82 8.233858e+54
## no_interaction_ML_pid 15210.94 15254.72 0.000000e+00
##
## $predicted_differences
##    0%   25%   50%   75%  100%
## 0.000 0.162 0.369 0.621 1.991
```

*Model summary*

```
## Linear mixed model fit by maximum likelihood . t-tests use Satterthwait
e's
##   method [lmerModLmerTest]
## Formula: outcome ~ decomp + intensity + condition + (1 | pid/muid)
##    Data: d
##
##      AIC      BIC   logLik deviance df.resid
##  14951.8  15001.8  -7467.9  14935.8     3832
##
## Scaled residuals:
##     Min      1Q  Median      3Q     Max
## -8.2392 -0.5476  0.0283  0.6115  3.8619
##
## Random effects:
##  Groups   Name        Variance Std.Dev.
##  muid:pid (Intercept) 0.5352   0.7316
##  pid      (Intercept) 2.0832   1.4433
##  Residual             2.5053   1.5828
## Number of obs: 3840, groups:  muid:pid, 432; pid, 17
##
## Fixed effects:
##                Estimate Std. Error         df t value Pr(>|t|)
## (Intercept)    12.30041    0.35755   17.69754  34.402  < 2e-16 ***
## decompMUedit   -0.05739    0.05479 3728.58958  -1.047    0.295
## intensity2      2.70995    0.05266 3584.95047  51.458  < 2e-16 ***
## condition2     -0.97023    0.06257 3399.56280 -15.507  < 2e-16 ***
## condition3      0.37516    0.06257 3399.56280   5.996 2.23e-09 ***
## ---
## Signif. codes:  0 '***' 0.001 '**' 0.01 '*' 0.05 '.' 0.1 ' ' 1
##
## Correlation of Fixed Effects:
##             (Intr) dcmpMU intns2 cndtn2
## decompMUedt -0.096
## intensity2  -0.060 -0.038
## condition2  -0.087  0.000  0.000
## condition3  -0.087  0.000  0.000  0.500
```

*Model- & sandwich-based standard errors and p-values*

The full cluster-robust variance-covariance matrix was calculated, which includes the variances (SEs) and covariances (how much the estimates correlate with each other):

```
## Model- & sandwich-based standard errors and p-values

## Click on the arrow for sandwich-based p-values.

## # A tibble: 5 × 6
##   term         estimate `Model-based s.e.` `Model-based p-value` `Sandw
ich s.e.`
##   <chr>           <dbl>              <dbl>                 <dbl>
<dbl>
## 1 (Intercept)   12.3               0.358             2.91e-226
0.342
## 2 decompMUedit  -0.0574            0.0548            2.95e-  1
0.0696
## 3 intensity2     2.71              0.0527            0
0.158
## 4 condition2    -0.970             0.0626            1.19e- 52
0.201
## 5 condition3     0.375             0.0626            2.21e-  9
0.152
##   `Sandwich p-value`
##                <dbl>
## 1          9.94e-17
## 2          4.23e- 1
## 3          7.93e-11
## 4          2.56e- 4
## 5          2.68e- 2
```

*Condition p-value*

The Wald statistic testing for an effect of condition is:

```
##  test Fstat df_num df_denom     p_val sig
##   HTZ  15.3      2     13.3 0.0003506 ***
```

*Post-hoc comparisons*

Estimated marginal means (EMMs) for the fixed-effects model with no interaction, while explicitly using cluster-robust standard errors (which accounts for non-independent data or heteroskedasticity), are reported below. This approach was used to ensure that the confidence intervals and subsequent p-values for the EMMs are more accurate and reliable.

```
##  intensity condition emmean    SE   df lower.CL upper.CL
##  1         1           12.3 0.337 17.4     11.6     13.0
##  2         1           15.0 0.437 17.5     14.1     15.9
##  1         2           11.3 0.368 17.4     10.5     12.1
##  2         2           14.0 0.430 17.5     13.1     14.9
##  1         3           12.6 0.354 17.4     11.9     13.4
##  2         3           15.4 0.440 17.5     14.4     16.3
##
## Results are averaged over the levels of: decomp
## Degrees-of-freedom method: satterthwaite
## Confidence level used: 0.95
```

```
## condition = 1:
##  contrast                 estimate    SE   df t.ratio p.value
##  intensity1 - intensity2    -2.71 0.158 3585 -17.152  <.0001
##
## condition = 2:
##  contrast                 estimate    SE   df t.ratio p.value
##  intensity1 - intensity2    -2.71 0.158 3585 -17.152  <.0001
##
## condition = 3:
##  contrast                 estimate    SE   df t.ratio p.value
##  intensity1 - intensity2    -2.71 0.158 3585 -17.152  <.0001
##
## Results are averaged over the levels of: decomp
## Degrees-of-freedom method: satterthwaite
```

```
## intensity = 1:
##  contrast                 estimate    SE   df t.ratio p.value
##  condition1 - condition2    0.970 0.201 3400   4.819  <.0001
##  condition1 - condition3   -0.375 0.152 3400  -2.467  0.0137
##  condition2 - condition3   -1.345 0.236 3400  -5.702  <.0001
##
## intensity = 2:
##  contrast                 estimate    SE   df t.ratio p.value
##  condition1 - condition2    0.970 0.201 3400   4.819  <.0001
##  condition1 - condition3   -0.375 0.152 3400  -2.467  0.0137
##  condition2 - condition3   -1.345 0.236 3400  -5.702  <.0001
##
## Results are averaged over the levels of: decomp
## Degrees-of-freedom method: satterthwaite
## P value adjustment: holm method for 3 tests
```

*Fixed-effects model with interaction*

This fixed-effects model fits decomp, intensity, condition and the interaction term intensity:condition with random intercepts and fixed slopes. The fixed-effects model with an intensity:condition interaction resulted in a better fit:

```
## $statistics
##                         aic      bic bayes.factor
## no_interaction_ML 14951.80 15001.82            0
## interaction_ML    14899.38 14961.91    464783854
##
## $predicted_differences
##    0%   25%   50%   75%  100%
## 0.000 0.034 0.205 0.237 0.292
```

*Model summary*

```
## Linear mixed model fit by maximum likelihood . t-tests use Satterthwait
e's
##   method [lmerModLmerTest]
## Formula: outcome ~ decomp + intensity * condition + (1 | pid/muid)
##    Data: d
##
```

```
##      AIC      BIC   logLik deviance df.resid
##  14899.4  14961.9  -7439.7  14879.4     3830
##
## Scaled residuals:
##     Min      1Q  Median      3Q     Max
## -8.2787 -0.5572  0.0257  0.6061  3.7193
##
## Random effects:
##  Groups   Name        Variance Std.Dev.
##  muid:pid (Intercept) 0.5408   0.7354
##  pid      (Intercept) 2.0828   1.4432
##  Residual             2.4641   1.5697
## Number of obs: 3840, groups:  muid:pid, 432; pid, 17
##
## Fixed effects:
##                           Estimate Std. Error         df t value Pr(>|t|)
## (Intercept)               12.49860    0.35895   17.98182  34.820  < 2e-16
***
## decompMUedit              -0.05747    0.05437 3725.65339  -1.057  0.29055
## intensity2                 2.27363    0.08891 3466.55848  25.572  < 2e-16
***
## condition2                -1.14040    0.08403 3398.97471 -13.572  < 2e-16
***
## condition3                -0.04943    0.08403 3398.97471  -0.588  0.55641
## intensity2:condition2      0.37425    0.12461 3398.97472   3.003  0.00269
**
## intensity2:condition3      0.93379    0.12461 3398.97472   7.494  8.5e-14
***
## ---
## Signif. codes:  0 '***' 0.001 '**' 0.01 '*' 0.05 '.' 0.1 ' ' 1
##
## Correlation of Fixed Effects:
##             (Intr) dcmpMU intns2 cndtn2 cndtn3 int2:2
## decompMUedt -0.095
## intensity2  -0.109 -0.022
## condition2  -0.117  0.000  0.473
## condition3  -0.117  0.000  0.473  0.500
## intnsty2:c2  0.079  0.000 -0.701 -0.674 -0.337
## intnsty2:c3  0.079  0.000 -0.701 -0.337 -0.674  0.500
```

*Model- & sandwich-based standard errors and p-values*

```
## Model- & sandwich-based standard errors and p-values

## Click on the arrow for sandwich-based p-values.

## # A tibble: 7 × 6
##   term                  estimate `Model-based s.e.` `Model-based p-valu
e`
##   <chr>                    <dbl>              <dbl>                 <db
l>
## 1 (Intercept)            12.5               0.359                4.84e-2
31
## 2 decompMUedit           -0.0575            0.0544               2.91e-
1
## 3 intensity2              2.27              0.0889               2.67e-1
33
## 4 condition2             -1.14              0.0840               5.14e-
41
## 5 condition3             -0.0494            0.0840               5.56e-
1
## 6 intensity2:condition2   0.374             0.125                2.69e-
3
## 7 intensity2:condition3   0.934             0.125                8.27e-
14
##   `Sandwich s.e.` `Sandwich p-value`
##             <dbl>              <dbl>
## 1          0.352            1.23e-16
## 2          0.0697           4.23e- 1
## 3          0.249            2.48e- 7
## 4          0.230            1.96e- 4
## 5          0.194            8.03e- 1
## 6          0.347            2.99e- 1
## 7          0.251            2.22e- 3
```

*Interaction, intensity & condition p-values*

The Wald statistics testing for an effect of the intensity:condition interaction term, for an effect of the intensity term and the intensity:condition interaction term, and for an effect of the condition term and the intensity:condition interaction term are:

```
##  test Fstat df_num df_denom     p_val sig
##   HTZ  12.7      2     13.3 0.0008294 ***

##  test Fstat df_num df_denom   p_val sig
##   HTZ  98.4      3     12.2 < 1e-04 ***

##  test Fstat df_num df_denom     p_val sig
##   HTZ  15.6      4     11.3 0.0001494 ***
```

*Post-hoc comparisons*

Estimated marginal means (EMMs) for the interaction between intensity and condition for the fixed-effects model, while explicitly using cluster-robust standard errors (which accounts for non-independent data or heteroskedasticity), are reported below. This approach was used to ensure that the confidence intervals and subsequent p-values for the EMMs are more accurate and reliable.

```
##  intensity condition emmean    SE   df lower.CL upper.CL
##  1         1           12.5 0.351 17.7     11.7     13.2
##  2         1           14.7 0.446 17.9     13.8     15.7
##  1         2           11.3 0.362 17.7     10.6     12.1
##  2         2           14.0 0.455 17.9     13.0     14.9
##  1         3           12.4 0.364 17.7     11.7     13.2
##  2         3           15.6 0.438 17.9     14.7     16.5
##
## Results are averaged over the levels of: decomp
## Degrees-of-freedom method: satterthwaite
## Confidence level used: 0.95

## condition = 1:
##  contrast                 estimate    SE   df t.ratio p.value
##  intensity1 - intensity2    -2.27 0.249 3467  -9.144  <.0001
##
## condition = 2:
##  contrast                 estimate    SE   df t.ratio p.value
##  intensity1 - intensity2    -2.65 0.236 3467 -11.205  <.0001
##
## condition = 3:
##  contrast                 estimate    SE   df t.ratio p.value
##  intensity1 - intensity2    -3.21 0.176 3467 -18.189  <.0001
##
## Results are averaged over the levels of: decomp
## Degrees-of-freedom method: satterthwaite

## intensity = 1:
##  contrast                 estimate    SE   df t.ratio p.value
##  condition1 - condition2   1.1404 0.230 3399   4.959  <.0001
##  condition1 - condition3   0.0494 0.194 3399   0.254  0.7992
##  condition2 - condition3  -1.0910 0.254 3399  -4.300  <.0001
##
## intensity = 2:
##  contrast                 estimate    SE   df t.ratio p.value
##  condition1 - condition2   0.7662 0.303 3399   2.530  0.0114
##  condition1 - condition3  -0.8844 0.205 3399  -4.312  <.0001
##  condition2 - condition3  -1.6505 0.260 3399  -6.359  <.0001
##
## Results are averaged over the levels of: decomp
## Degrees-of-freedom method: satterthwaite
## P value adjustment: holm method for 3 tests
```

*Random-effects model with interaction*

This random-effects model fits decomp, intensity, condition and the interaction term intensity:condition with random intercepts and random slopes. The random-effects model with an intensity:condition interaction resulted in a better fit:

```
## $statistics
##                          aic      bic bayes.factor
## interaction_ML        14899.38 14961.91            0
## interaction_ML_random 12389.57 12789.77          Inf
##
## $predicted_differences
##    0%   25%   50%   75%  100%
## 0.000 0.353 0.773 1.414 5.417
```

*Model summary*

```
## Linear mixed model fit by maximum likelihood . t-tests use Satterthwait
e's
##   method [lmerModLmerTest]
## Formula: outcome ~ decomp + intensity * condition + (decomp + intensity
*
##     condition | pid/muid)
##    Data: d
##
##      AIC      BIC   logLik deviance df.resid
##  12389.6  12789.8  -6130.8  12261.6     3776
##
## Scaled residuals:
##      Min       1Q   Median       3Q      Max
## -11.2798  -0.4274   0.0124   0.4772   4.5406
##
## Random effects:
##  Groups   Name                  Variance Std.Dev. Corr
##  muid:pid (Intercept)           1.504330 1.22651
##           decompMUedit          2.850828 1.68844  -0.68
##           intensity2            3.207226 1.79087  -0.32 -0.16
##           condition2            0.012766 0.11299   0.86 -0.24 -0.37
##           condition3            0.004523 0.06726   0.98 -0.52 -0.35  0.
95
##           intensity2:condition2 0.042466 0.20607  -0.62  0.11  0.94 -0.
61 -0.64
##           intensity2:condition3 0.004587 0.06773  -0.97  0.76  0.07 -0.
80 -0.94
##  pid      (Intercept)           1.968890 1.40317
##           decompMUedit          0.022747 0.15082   0.11
##           intensity2            0.531316 0.72891   0.05 -0.38
##           condition2            0.942116 0.97063  -0.34 -0.36  0.33
##           condition3            0.520861 0.72171  -0.22 -0.60  0.32  0.
14
##           intensity2:condition2 1.808252 1.34471   0.40  0.29 -0.81 -0.
69 -0.32
##           intensity2:condition3 1.005440 1.00272   0.35  0.06 -0.72 -0.
47 -0.39
##  Residual                       0.670356 0.81875
##
##
##
##
##
```

```
##
##
##   0.41
##
##
##
##
##
##
##   0.79
##
## Number of obs: 3840, groups:  muid:pid, 432; pid, 17
##
## Fixed effects:
##                         Estimate Std. Error        df t value Pr(>|t|)
## (Intercept)            12.464182   0.350039 16.599283  35.608  < 2e-16 *
**
## decompMUedit           -0.013230   0.103706 48.325978  -0.128  0.89902
## intensity2              2.318391   0.206591 17.434180  11.222 2.11e-09 *
**
## condition2             -1.253109   0.240094 16.729353  -5.219 7.31e-05 *
**
## condition3             -0.008008   0.181170 16.849509  -0.044  0.96526
## intensity2:condition2  0.416430   0.333747 16.752620   1.248  0.22928
## intensity2:condition3  0.971166   0.252993 16.406152   3.839  0.00139 *
*
## ---
## Signif. codes:  0 '***' 0.001 '**' 0.01 '*' 0.05 '.' 0.1 ' ' 1
##
## Correlation of Fixed Effects:
##             (Intr) dcmpMU intns2 cndtn2 cndtn3 int2:2
## decompMUedt -0.114
## intensity2   0.001 -0.174
## condition2  -0.337 -0.133  0.299
## condition3  -0.222 -0.214  0.292  0.160
## intnsty2:c2  0.389  0.105 -0.698 -0.688 -0.320
## intnsty2:c3  0.337  0.029 -0.637 -0.457 -0.414  0.775
## optimizer (nloptwrap) convergence code: 0 (OK)
## boundary (singular) fit: see help('isSingular')
```

*Model- & sandwich-based standard errors and p-values*

```
## Model- & sandwich-based standard errors and p-values

## Click on the arrow for sandwich-based p-values.

## # A tibble: 7 × 6
##   term                  estimate `Model-based s.e.` `Model-based p-valu
e`
##   <chr>                    <dbl>              <dbl>                 <db
l>
## 1 (Intercept)          12.5                   0.350             1.05e-2
39
## 2 decompMUedit         -0.0132                0.104             8.98e-
1
## 3 intensity2            2.32                  0.207             9.02e-
29
## 4 condition2           -1.25                  0.240             1.89e-
7
## 5 condition3           -0.00801               0.181             9.65e-
1
## 6 intensity2:condition2  0.416                0.334             2.12e-
1
## 7 intensity2:condition3  0.971                0.253             1.26e-
4
##   `Sandwich s.e.` `Sandwich p-value`
##             <dbl>              <dbl>
## 1          0.356           1.55e-16
## 2          0.0802          8.71e- 1
## 3          0.212           8.66e- 9
## 4          0.247           1.15e- 4
## 5          0.187           9.66e- 1
## 6          0.344           2.44e- 1
## 7          0.261           1.84e- 3
```

*Interaction, intensity & condition p-values*

The Wald statistics testing for an effect of the intensity:condition interaction term, for an effect of the intensity term and the intensity:condition interaction term, and for an effect of the condition term and the intensity:condition interaction term are:

```
##  test Fstat df_num df_denom    p_val sig
##   HTZ  9.81      2       15 0.001893  **

##  test Fstat df_num df_denom   p_val sig
##   HTZ   100      3     13.8 < 1e-04 ***

##  test Fstat df_num df_denom     p_val sig
##   HTZ  13.6      4       13 0.0001409 ***
```

*Post-hoc comparisons*

Estimated marginal means (EMMs) for the interaction between intensity and condition for the random-effects model, while explicitly using cluster-robust standard errors (which accounts for non-independent data or heteroskedasticity), are reported below. This approach was used to ensure that the confidence intervals and subsequent p-values for the EMMs are more accurate and reliable.

```
##  intensity condition emmean    SE   df lower.CL upper.CL
##  1         1           12.5 0.358 16.7     11.7     13.2
##  2         1           14.8 0.413 16.8     13.9     15.6
##  1         2           11.2 0.355 16.9     10.5     12.0
##  2         2           13.9 0.445 16.6     13.0     14.9
##  1         3           12.4 0.359 16.7     11.7     13.2
##  2         3           15.7 0.464 16.5     14.8     16.7
##
## Results are averaged over the levels of: decomp
## Degrees-of-freedom method: satterthwaite
## Confidence level used: 0.95
```

```
## condition = 1:
##  contrast                 estimate    SE   df t.ratio p.value
##  intensity1 - intensity2     -2.32 0.212 17.4 -10.950  <.0001
##
## condition = 2:
##  contrast                 estimate    SE   df t.ratio p.value
##  intensity1 - intensity2     -2.73 0.246 14.5 -11.104  <.0001
##
## condition = 3:
##  contrast                 estimate    SE   df t.ratio p.value
##  intensity1 - intensity2     -3.29 0.205 14.0 -16.072  <.0001
##
## Results are averaged over the levels of: decomp
## Degrees-of-freedom method: satterthwaite
```

```
## intensity = 1:
##  contrast                 estimate    SE   df t.ratio p.value
##  condition1 - condition2  1.25311 0.247 16.7   5.066  0.0003
##  condition1 - condition3  0.00801 0.187 16.9   0.043  0.9663
##  condition2 - condition3 -1.24510 0.285 16.8  -4.367  0.0009
##
## intensity = 2:
##  contrast                 estimate    SE   df t.ratio p.value
##  condition1 - condition2  0.83668 0.250 17.1   3.350  0.0038
##  condition1 - condition3 -0.96316 0.250 16.2  -3.852  0.0028
##  condition2 - condition3 -1.79984 0.267 16.8  -6.745  <.0001
##
## Results are averaged over the levels of: decomp
## Degrees-of-freedom method: satterthwaite
## P value adjustment: holm method for 3 tests
```

### Coefficient of variation in motor unit discharge rate model fitting and results

Fixed effects were fitted using a linear mixed-effects model with motor units as random effects nested under the participant id. Model-based standard errors and p-values are presented. Sandwich variance estimation-based standard errors and p-values were also produced; these do not assume normality of the data. A Wald statistic based on the sandwich variance estimate was also produced to test the effect of factor condition.

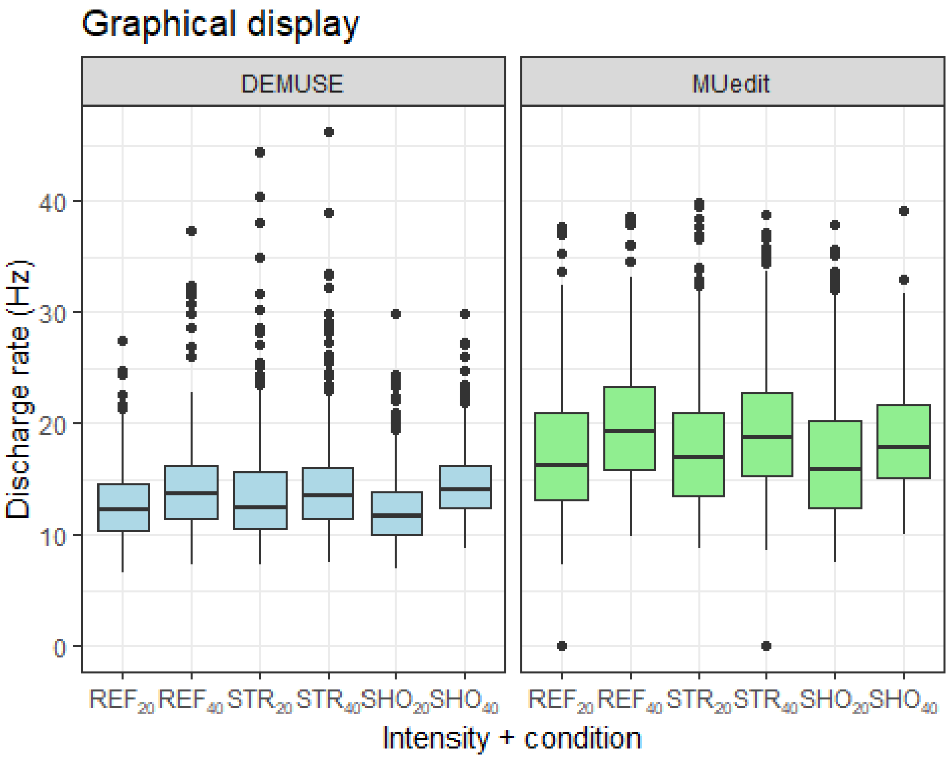


*Fixed-effects model with no interaction and with/without random motor unit effects nested under random participant effects*

This fixed-effects model fits the factors decomp, intensity, and condition with random intercepts and fixed slopes. We first compared fitting the model by REML vs. ML, and ML resulted in a better fit with random motor unit effects nested under random participant effects:

```
## $statistics
##                          aic      bic bayes.factor
## no_interaction_REML 23099.23 23149.26        0.030
## no_interaction_ML   23092.22 23142.24       33.326
##
## $predicted_differences
##    0%   25%   50%   75%  100%
## 0.000 0.001 0.003 0.004 0.012
```

```
## $statistics
##                              aic      bic bayes.factor
## no_interaction_ML     23092.22 23142.24 5.252722e+23
## no_interaction_ML_pid 23207.71 23251.48 0.000000e+00
##
## $predicted_differences
##    0%   25%   50%   75%  100%
## 0.001 0.410 0.844 1.333 4.039
```

*Model summary*

```
## Linear mixed model fit by maximum likelihood . t-tests use Satterthwait
e's
##   method [lmerModLmerTest]
## Formula: outcome ~ decomp + intensity + condition + (1 | pid/muid)
##    Data: d
##
##      AIC      BIC   logLik deviance df.resid
##  23092.2  23142.2 -11538.1  23076.2     3832
##
## Scaled residuals:
##     Min      1Q  Median      3Q     Max
## -4.7329 -0.6310 -0.1518  0.4158  6.0168
##
## Random effects:
##  Groups   Name        Variance Std.Dev.
##  muid:pid (Intercept)  2.876   1.696
##  pid      (Intercept)  2.660   1.631
##  Residual             21.665   4.655
## Number of obs: 3840, groups:  muid:pid, 432; pid, 17
##
## Fixed effects:
##               Estimate Std. Error        df t value Pr(>|t|)
## (Intercept)    13.4226     0.4436   22.5146  30.257  < 2e-16 ***
## decompMUedit    4.1896     0.1597 3766.7249  26.237  < 2e-16 ***
## intensity2      2.0864     0.1541 3593.1192  13.536  < 2e-16 ***
## condition2      0.4043     0.1840 3364.0185   2.197  0.02807 *
## condition3     -0.6042     0.1840 3364.0185  -3.284  0.00103 **
## ---
## Signif. codes:  0 '***' 0.001 '**' 0.01 '*' 0.05 '.' 0.1 ' ' 1
##
## Correlation of Fixed Effects:
##             (Intr) dcmpMU intns2 cndtn2
## decompMUedt -0.223
## intensity2  -0.145 -0.034
## condition2  -0.207  0.000  0.000
## condition3  -0.207  0.000  0.000  0.500
```

*Model- & sandwich-based standard errors and p-values*

The full cluster-robust variance-covariance matrix was calculated, which includes the variances (SEs) and covariances (how much the estimates correlate with each other):

```
## Model- & sandwich-based standard errors and p-values

## Click on the arrow for sandwich-based p-values.

## # A tibble: 5 × 6
##   term         estimate `Model-based s.e.` `Model-based p-value` `Sandw
ich s.e.`
##   <chr>           <dbl>              <dbl>                 <dbl>
<dbl>
## 1 (Intercept)    13.4                0.444             1.61e-180
0.598
## 2 decompMUedit    4.19               0.160             1.11e-139
0.327
## 3 intensity2      2.09               0.154             8.23e- 41
0.610
## 4 condition2      0.404              0.184             2.81e-  2
0.399
## 5 condition3     -0.604              0.184             1.03e-  3
0.340
##   `Sandwich p-value`
##                <dbl>
## 1           1.78e-13
## 2           2.58e- 9
## 3           4.09e- 3
## 4           3.27e- 1
## 5           9.70e- 2
```

*Fixed-effects model with interaction*

This fixed-effects model fits decomp, intensity, condition and the interaction term intensity:condition with random intercepts and fixed slopes. The fixed-effects model with an intensity:condition interaction did not necessarily result in a better fit:

```
## $statistics
##                        aic      bic bayes.factor
## no_interaction_ML 23092.22 23142.24      327.993
## interaction_ML    23091.30 23153.83        0.003
##
## $predicted_differences
##    0%   25%   50%   75%  100%
## 0.021 0.033 0.173 0.204 0.242
```

*Random-effects model with no interaction*

This random-effects model fits decomp, intensity and condition with random intercepts and random slopes. The random-effects model with no interaction resulted in a better fit:

```
## $statistics
##                               aic      bic bayes.factor
## no_interaction_ML        23092.22 23142.24 0.000000e+00
## no_interaction_ML_random 22645.39 22870.51 1.015758e+59
##
```

```
## $predicted_differences
##    0%   25%   50%   75%  100%
## 0.001 0.602 1.304 2.271 7.733
```

*Model summary*

```
## Linear mixed model fit by maximum likelihood . t-tests use Satterthwait
e's
##   method [lmerModLmerTest]
## Formula: outcome ~ decomp + intensity + condition + (decomp + intensity
+
##     condition | pid/muid)
##    Data: d
##
##      AIC      BIC   logLik deviance df.resid
##  22645.4  22870.5 -11286.7  22573.4     3804
##
## Scaled residuals:
##     Min      1Q  Median      3Q     Max
## -4.2906 -0.5471 -0.1465  0.3404  5.3293
##
## Random effects:
##  Groups   Name         Variance Std.Dev. Corr
##  muid:pid (Intercept)   4.2028  2.0501
##           decompMUedit  9.4817  3.0792   -0.45
##           intensity2    6.4933  2.5482   -0.39 -0.20
##           condition2    0.5848  0.7647    0.62 -0.24  0.39
##           condition3    0.2170  0.4658   -0.09 -0.07 -0.73 -0.84
##  pid      (Intercept)   5.0376  2.2445
##           decompMUedit  0.7682  0.8765   -0.36
##           intensity2    4.2165  2.0534   -0.15  0.06
##           condition2    2.4339  1.5601   -0.48 -0.12 -0.10
##           condition3    1.1876  1.0898   -0.73  0.53 -0.37  0.44
##  Residual              15.4904  3.9358
## Number of obs: 3840, groups:  muid:pid, 432; pid, 17
##
## Fixed effects:
##              Estimate Std. Error      df t value Pr(>|t|)
## (Intercept)   13.3353     0.5797 16.7459  23.004 4.16e-14 ***
## decompMUedit   4.3838     0.3038 18.2660  14.430 1.97e-11 ***
## intensity2     1.8632     0.5371 17.1559   3.469   0.0029 **
## condition2     0.5112     0.4142 16.6244   1.234   0.2343
## condition3    -0.5803     0.3103 17.3466  -1.870   0.0785 .
## ---
## Signif. codes:  0 '***' 0.001 '**' 0.01 '*' 0.05 '.' 0.1 ' ' 1
##
## Correlation of Fixed Effects:
##             (Intr) dcmpMU intns2 cndtn2
## decompMUedt -0.366
## intensity2  -0.179  0.011
## condition2  -0.457 -0.092 -0.071
## condition3  -0.659  0.323 -0.313  0.438
## optimizer (nloptwrap) convergence code: 0 (OK)
## boundary (singular) fit: see help('isSingular')
```

*Model- & sandwich-based standard errors and p-values*

```
## Model- & sandwich-based standard errors and p-values

## Click on the arrow for sandwich-based p-values.

## # A tibble: 5 × 6
##   term        estimate `Model-based s.e.` `Model-based p-value` `Sandw
ich s.e.`
##   <chr>          <dbl>              <dbl>                 <dbl>
<dbl>
## 1 (Intercept)    13.3               0.580             9.56e-110
0.598
## 2 decompMUedit    4.38              0.304             5.36e- 46
0.313
## 3 intensity2      1.86              0.537             5.28e-  4
0.553
## 4 condition2      0.511             0.414             2.17e-  1
0.427
## 5 condition3     -0.580             0.310             6.16e-  2
0.320
##   `Sandwich p-value`
##                <dbl>
## 1           1.83e-13
## 2           3.24e-10
## 3           3.94e- 3
## 4           2.49e- 1
## 5           8.89e- 2
```

*Estimated marginal means*

Estimated marginal means (EMMs) for the random-effects model with no interaction are reported below.

```
##  decomp emmean    SE   df lower.CL upper.CL
##  DEMUSE   14.2 0.484 17.1     13.2     15.3
##  MUedit   18.6 0.458 17.0     17.7     19.6
## 
## Results are averaged over the levels of: intensity, condition
## Degrees-of-freedom method: satterthwaite
## Confidence level used: 0.95

##  intensity emmean    SE   df lower.CL upper.CL
##  1           15.5 0.451 16.5     14.6     16.5
##  2           17.4 0.587 17.2     16.1     18.6
## 
## Results are averaged over the levels of: decomp, condition
## Degrees-of-freedom method: satterthwaite
## Confidence level used: 0.95

##  condition emmean    SE   df lower.CL upper.CL
##  1           16.5 0.576 16.9     15.2     17.7
##  2           17.0 0.502 17.4     15.9     18.0
##  3           15.9 0.398 16.6     15.0     16.7
## 
```

```
## Results are averaged over the levels of: decomp, intensity
## Degrees-of-freedom method: satterthwaite
## Confidence level used: 0.95
```

### Rate of agreement model fitting and results

*Manually edited DEMUSE decompositions versus MUedit*

Fixed effects were fitted using a linear mixed-effects model with and without motor units as random effects nested under the participant id. Model-based standard errors and p-values are presented. Sandwich variance estimation-based standard errors and p-values were also produced; these do not assume normality of the data.

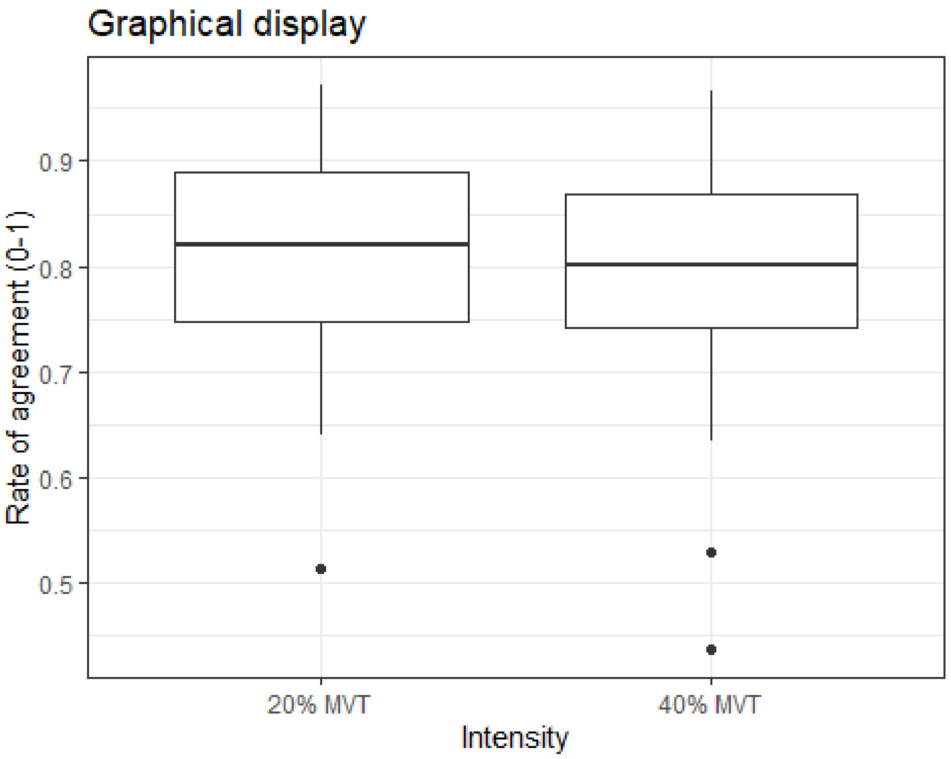


*Fixed-effects model with/without random motor unit effects nested under random participant effects*

This fixed-effects model fits the factor intensity with random intercepts and fixed slopes. We first compared fitting the model by REML vs. ML, and ML resulted in a better fit without random motor unit effects nested under random participant effects:

```
## $statistics
##                          aic      bic bayes.factor
## no_interaction_REML -972.34 -951.727        0.000
## no_interaction_ML   -989.49 -968.877     5296.999
##
## $predicted_differences
##    0%   25%   50%   75%  100%
## 0.000 0.000 0.001 0.001 0.002

## $statistics
##                            aic      bic bayes.factor
## no_interaction_ML     -989.49 -968.877        0.047
## no_interaction_ML_pid -991.49 -975.000       21.354
```

```
##


## $predicted_differences
##   0%  25%  50%  75% 100%
##    0    0    0    0    0
```

*Model summary*

```
## Linear mixed model fit by maximum likelihood . t-tests use Satterthwait
e's
##   method [lmerModLmerTest]
## Formula: outcome ~ intensity + (1 | pid)
##    Data: d
##
##      AIC      BIC   logLik deviance df.resid
##   -991.5   -975.0    499.7   -999.5      452
##
## Scaled residuals:
##     Min      1Q  Median      3Q     Max
## -4.5201 -0.7892 -0.0056  0.8326  2.0173
##
## Random effects:
##  Groups   Name        Variance  Std.Dev.
##  pid      (Intercept) 8.733e-06 0.002955
##  Residual             6.532e-03 0.080820
## Number of obs: 456, groups:  pid, 17
##
## Fixed effects:
##              Estimate Std. Error         df t value Pr(>|t|)
## (Intercept)   0.820059   0.005089  42.474173 161.143   <2e-16 ***
## intensity2   -0.017544   0.007638 450.326724  -2.297   0.0221 *
## ---
## Signif. codes:  0 '***' 0.001 '**' 0.01 '*' 0.05 '.' 0.1 ' ' 1
##
## Correlation of Fixed Effects:
##            (Intr)
## intensity2 -0.651
```

*Model- & sandwich-based standard errors and p-values*

The full cluster-robust variance-covariance matrix was calculated, which includes the variances (SEs) and covariances (how much the estimates correlate with each other):

```
## Model- & sandwich-based standard errors and p-values

## Click on the arrow for sandwich-based p-values.

## # A tibble: 2 × 6
##   term        estimate `Model-based s.e.` `Model-based p-value` `Sandwi
ch s.e.`
##   <chr>          <dbl>              <dbl>                 <dbl>
<dbl>
## 1 (Intercept)   0.820            0.00509                0
0.00600
## 2 intensity2   -0.0175           0.00764                0.0221
0.00663
##   `Sandwich p-value`
##              <dbl>
## 1          2.26e-23
## 2          1.93e- 2
```

*Estimated marginal means*

Estimated marginal means (EMMs) for the fixed-effects model, while explicitly using cluster-robust standard errors (which accounts for non-independent data or heteroskedasticity), are reported below. This approach was used to ensure that the confidence intervals and subsequent p-values for the EMMs are more accurate and reliable.

```
##  intensity emmean      SE   df lower.CL upper.CL
##  1          0.820 0.00600 42.5    0.808    0.832
##  2          0.803 0.00383 65.2    0.795    0.810
##
## Degrees-of-freedom method: satterthwaite
## Confidence level used: 0.95
```

*Random-effects model*

This random-effects model fits intensity with random intercepts and random slopes.  The fixed-effects model resulted in a better fit:

```
## $statistics
##                                   aic      bic bayes.factor
## no_interaction_ML_pid        -991.490 -975.000      315.846
## no_interaction_ML_pid_random -988.224 -963.489        0.003
##
## $predicted_differences
##    0%   25%   50%   75%  100%
## 0.000 0.000 0.001 0.005 0.011
```

*Unedited DEMUSE decompositions versus MUedit*

Fixed effects were fitted using a linear mixed-effects model with and without motor units as random effects nested under the participant id. Model-based standard errors and p-values are presented. Sandwich variance estimation-based standard errors and p-values were also produced; these do not assume normality of the data.

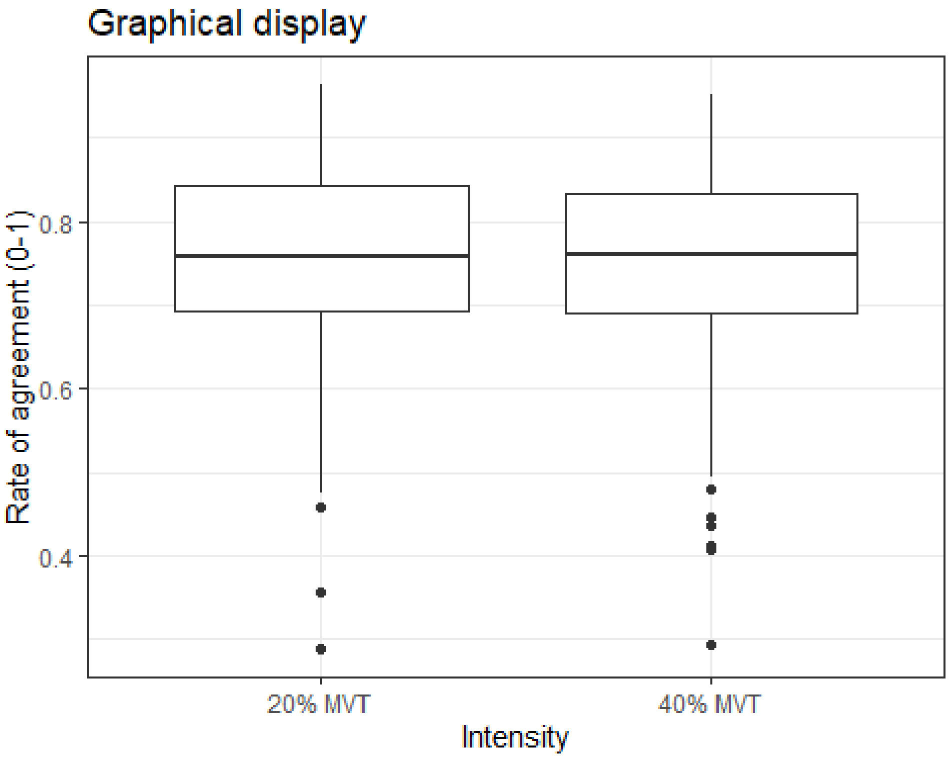


*Fixed-effects model with/without random motor unit effects nested under random participant effects*

This fixed-effects model fits the factor intensity with random intercepts and fixed slopes. We first compared fitting the model by REML vs. ML, and ML resulted in a better fit without random motor unit effects nested under random participant effects:

```
## $statistics
##                            aic      bic bayes.factor
## no_interaction_REML -745.475 -724.564        0.000
## no_interaction_ML   -761.219 -740.308     2622.514
##
## $predicted_differences
##    0%   25%   50%   75%  100%
## 0.000 0.000 0.000 0.001 0.003

## $statistics
##                              aic      bic bayes.factor
## no_interaction_ML     -761.219 -740.308        0.045
## no_interaction_ML_pid -763.219 -746.490       22.000
##
```

```
## $predicted_differences
##   0%  25%  50%  75% 100%
##    0    0    0    0    0
```

*Model summary*

```
## Linear mixed model fit by maximum likelihood . t-tests use Satterthwait
e's
##   method [lmerModLmerTest]
## Formula: outcome ~ intensity + (1 | pid)
##    Data: d
##
##      AIC      BIC   logLik deviance df.resid
##   -763.2   -746.5    385.6   -771.2      480
##
## Scaled residuals:
##     Min      1Q  Median      3Q     Max
## -4.2329 -0.5955  0.0637  0.7043  1.8751
##
## Random effects:
##  Groups   Name        Variance  Std.Dev.
##  pid      (Intercept) 0.0001787 0.01337
##  Residual             0.0117521 0.10841
## Number of obs: 484, groups:  pid, 17
##
## Fixed effects:
##               Estimate Std. Error        df t value Pr(>|t|)
## (Intercept)   0.75835    0.00738  33.85659 102.751   <2e-16 ***
## intensity2   -0.01014    0.00996 475.73147  -1.018    0.309
## ---
## Signif. codes:  0 '***' 0.001 '**' 0.01 '*' 0.05 '.' 0.1 ' ' 1
##
## Correlation of Fixed Effects:
##            (Intr)
## intensity2 -0.586
```

*Model- & sandwich-based standard errors and p-values*

The full cluster-robust variance-covariance matrix was calculated, which includes the variances (SEs) and covariances (how much the estimates correlate with each other):

```
## Model- & sandwich-based standard errors and p-values

## Click on the arrow for sandwich-based p-values.

## # A tibble: 2 × 6
##   term        estimate `Model-based s.e.` `Model-based p-value` `Sandwi
ch s.e.`
##   <chr>          <dbl>              <dbl>                 <dbl>
<dbl>
## 1 (Intercept)   0.758            0.00738                 0
0.00946
## 2 intensity2   -0.0101           0.00996                 0.309
0.0137
##   `Sandwich p-value`
##                <dbl>
## 1           7.15e-21
## 2           4.71e- 1
```

*Estimated marginal means*

Estimated marginal means (EMMs) for the fixed-effects model, while explicitly using cluster-robust standard errors (which accounts for non-independent data or heteroskedasticity), are reported below. This approach was used to ensure that the confidence intervals and subsequent p-values for the EMMs are more accurate and reliable.

```
##  intensity emmean      SE   df lower.CL upper.CL
##  1          0.758 0.00946 33.9    0.739    0.778
##  2          0.748 0.00879 48.5    0.731    0.766
## 
## Degrees-of-freedom method: satterthwaite 
## Confidence level used: 0.95
```

*Random-effects model*

This random-effects model fits intensity with random intercepts and random slopes. The fixed-effects model resulted in a better fit:

```
## $statistics
##                                   aic      bic bayes.factor
## no_interaction_ML_pid        -763.219 -746.490       86.440
## no_interaction_ML_pid_random -762.664 -737.571        0.012
## 
## $predicted_differences
##    0%   25%   50%   75%  100% 
## 0.001 0.005 0.008 0.012 0.024
```

### References

Avrillon S, Hug F, Baker SN, Gibbs C & Farina D (2024). Tutorial on MUedit: An open-source software for identifying and analysing the discharge timing of motor units from electromyographic signals. *J Electromyogr Kinesiol* **77,** 102886.

Del Vecchio A, Holobar A, Falla D, Felici F, Enoka RM & Farina D (2020). Tutorial: Analysis of motor unit discharge characteristics from high-density surface EMG signals. *J Electromyogr Kinesiol* **53,** 102426.

Holobar A, Minetto MA, Botter A, Negro F & Farina D (2010). Experimental analysis of accuracy in the identification of motor unit spike trains from high-density surface EMG. *IEEE Trans Neural Syst Rehabil Eng* **18,** 221–229.

Holobar A & Zazula D (2007*a*). Gradient convolution kernel compensation applied to surface electromyograms. In *Independent Component Analysis and Signal Separation*, ed. Davies ME, James CJ, Abdallah SA & Plumbley MD, Lecture Notes in Computer Science, pp. 617–624. Springer Berlin Heidelberg, Berlin, Heidelberg. Available at: http://link.springer.com/10.1007/978-3-540-74494-8_77 [Accessed August 14, 2025].

Holobar A & Zazula D (2007*b*). Multichannel blind source separation using convolution kernel compensation. *IEEE Trans Signal Process* **55,** 4487–4496.

Hug F, Avrillon S, Del Vecchio A, Casolo A, Ibanez J, Nuccio S, Rossato J, Holobar A & Farina D (2021). Analysis of motor unit spike trains estimated from high-density surface electromyography is highly reliable across operators. *J Electromyogr Kinesiol* **58,** 102548.

Murks N, Škarabot J, Kramberger M, Divjak M, Sedej G, Valenčič T, Connelly CD, Thomason H & Holobar A (2025). The efficiency of manual editing of high-density surface electromyogram decomposition depends on the recorded muscle and contraction level but less on the operator's experience. *IEEE Trans Neural Syst Rehabil Eng* **33,** 3367–3376.

Wade L, Needham L, Evans M, McGuigan P, Colyer S, Cosker D & Bilzon J (2023). Examination of 2D frontal and sagittal markerless motion capture: Implications for markerless applications ed. Gu Y. *PLoS ONE* **18,** e0293917.
